# Adaptive planning depth in human problem solving

**DOI:** 10.1101/2023.05.02.539099

**Authors:** Mattia Eluchans, Gian Luca Lancia, Antonella Maselli, Marco D’Alessando, Jeremy Gordon, Giovanni Pezzulo

## Abstract

We humans are capable of solving challenging planning problems, but the range of adaptive strategies that we use to address them are not yet fully characterized. Here, we designed a series of problem-solving tasks that require planning at different depths. After systematically comparing the performance of participants and planning models, we found that when facing problems that require planning to a certain number of subgoals (from 1 to 8), participants make an adaptive use of their cognitive resources – namely, they tend to select an initial plan having the minimum required depth, rather than selecting the same depth for all problems. These results support the view of problem solving as a bounded rational process, which adapts costly cognitive resources to task demands.

## Introduction

Since the early days of cognitive science, researchers have asked how we solve challenging problems that engage planning abilities, such as the Tower of Hanoi and Traveling Salesman as well as popular games such as chess or go (1–3). Most cognitive theories assume that planning requires a form of cognitive *tree search* over an internal model or mental map of the task (4–8). Following this view, several planning studies in humans and other animals used tree-like tasks, but mostly focused on simple (e.g., two-step) problems which could be searched exhaustively (9–12). It is still unclear how we solve more complex problems, which – given our limited resources – defy exhaustive search.

From a normative perspective, planning with limited resources could be described as a *bounded* rational processes, which balances the accuracy of the solution and the cognitive resources invested, e.g., memory and time (13–16). One way to lower cognitive resources is using *heuristics* to alleviate the burden of exhaustive search (17,18). For example, it has been proposed that people use a pruning heuristic during mental search: if they encounter a tree node that seems unpromising, they discard the whole branch of the tree (19–21). Other heuristics consist of sampling only a few promising routes, or many routes but only up to a certain depth (22,23). The tradition of ecological rationality highlights the use of various other smart, simple and fast heuristics that provide adaptive solutions to challenging problems (24,25). It also highlights *embodied* heuristics that exploit sensory and motor capabilities to facilitate effective decisions (26,27). Furthermore, it is possible to alleviate the burden of planning by using a hierarchical approach to split the problem into more manageable subproblems (28–32) or by interleaving planning and execution; for example, plan until a certain subgoal, then revise and complete the plan along the way, as one moves toward the chosen subgoal (17,18). Despite this progress, we still have incomplete knowledge of the (approximate) planning methods that humans and other animals might adopt during problem solving, as well as their neuronal underpinning (33–35).

Another stream of research explored the limitations (e.g., the maximum depth) of our planning abilities. Various studies have shown that with sufficient time, people are able to find near-optimal solutions to challenging problems, such as the Traveling Salesman, which requires finding the shortest possible closed path that connects a fixed number of “cities”. Different explanations of the Traveling Salesman are based on the idea that participants could solve complex problems by using local planning (36) or by avoiding planning ahead in depth and instead valuating the global perceptual properties of the problem to generate their solution (37). Other studies have tried to quantify planning depth in chess (38,39) and other games (40), see (41) for a recent review. Classical studies reported that top chess players can plan ahead (on average) a relatively small number of moves, between 3.6 and 5.4 and their maximum planning depth is between 6.8 and 9.1 moves (39,42,43) and a recent large-scale study reported that planning depth in games increases with expertise up to a ceiling at a similar level (44); but see (45) for evidence that chess grand masters can plan ahead (on average) 13.8 moves.

Planning – especially when done at greater depths – requires engaging significant cognitive resources and hence a key question is whether people make an adaptive or resource-rational (46,47) use of these resources. Deciding the appropriate planning depth for a particular problem can be seen as an instance of metaplanning: namely, the formation of a plan about how to approach the planning problem (16,48–51). One possibility is that people are sensitive to the planning demands of different problems and flexibly adapt their planning depth to the minimum depth necessary to solve them. An alternative possibility is that people are insensitive to planning demands and use a fixed planning depth to solve all the problems.

To disambiguate between these hypotheses, we asked participants to solve a series of planning problems that required finding a path to connect all the “gems” in a grid, without passing through the same node twice (though “backtracking” to un-select nodes was allowed). Participants solved the problems by “navigating” with a finger in a grid that was fully visible on their mobile phones. They had 60 seconds to solve each problem and earned more points if they solved it faster. Figure 1 shows an example problem, which requires finding a path from the home location (yellow node) through all the gems (red nodes). The six panels show six representative timesteps of the solution, with the azure line indicating the path taken (visible to the participant) and the small red dots showing the actual finger positions at different times (not visible to participants). In the example illustrated in Figure 1, the participants see on their mobile phone the configuration shown in Panel A. They first select an incorrect path towards the two gems to the right (Panel B), then they backtrack to the home location (Panels C-D), and finally select a correct path that connects all the gems (Panels E-F), solving the problem.

**Figure 1.**
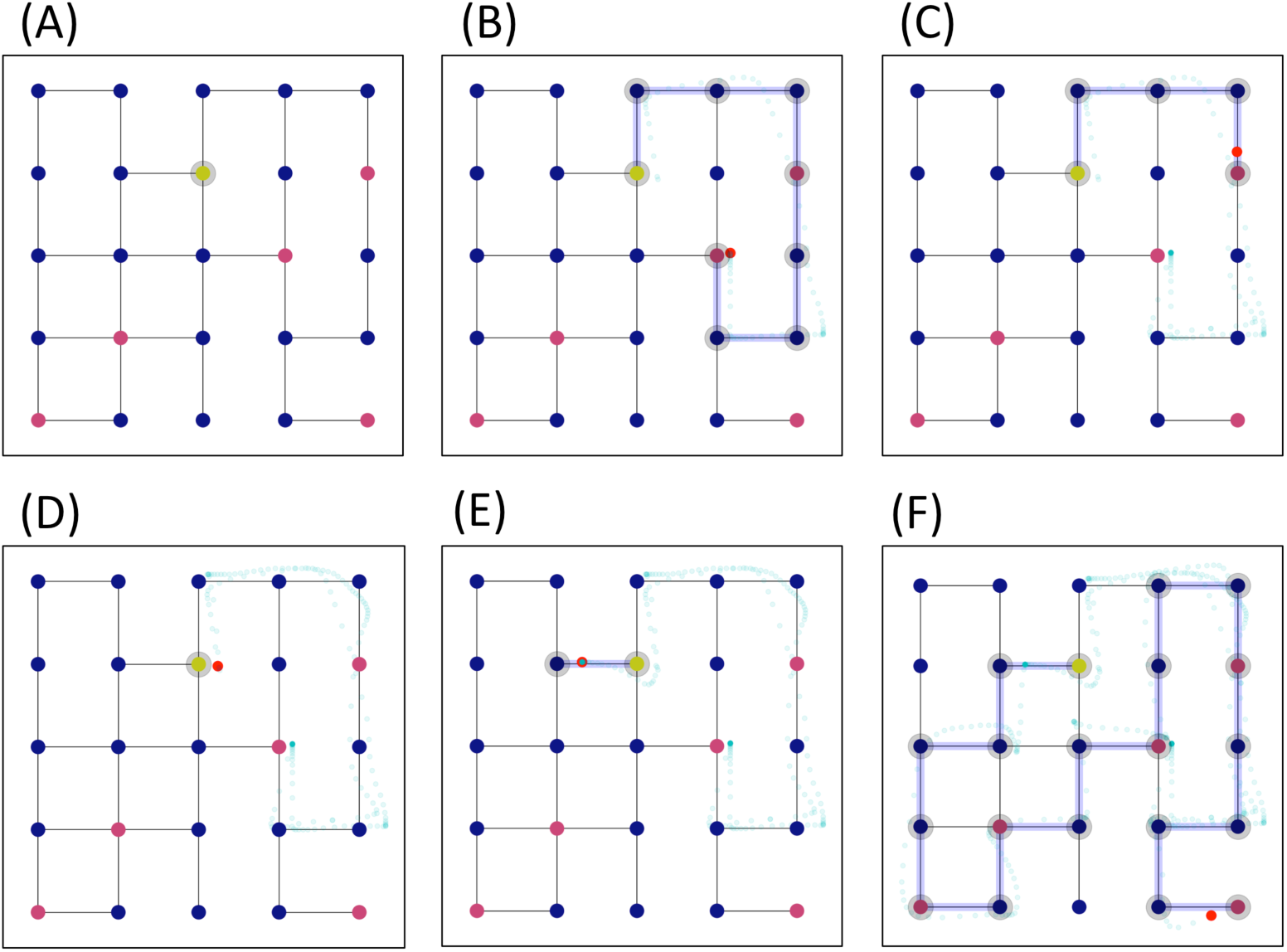
An example problem in the experiment. The problem requires finding a path in the grid that starts from the home location (yellow node) and collects all the “gems” (red nodes), without passing through the same node twice. Participants solved the problem by navigating with their finger on the (fully visible) grid, on their mobile phones. The figure shows six time-steps of the solution, with the azure path indicating the path taken by one of the participants, the azure dots the actual finger trajectory (sampled at 60Hz) and the small red dot the current finger position. Note that solving this particular problem requires planning 5 gems in advance.

**Figure 2.**
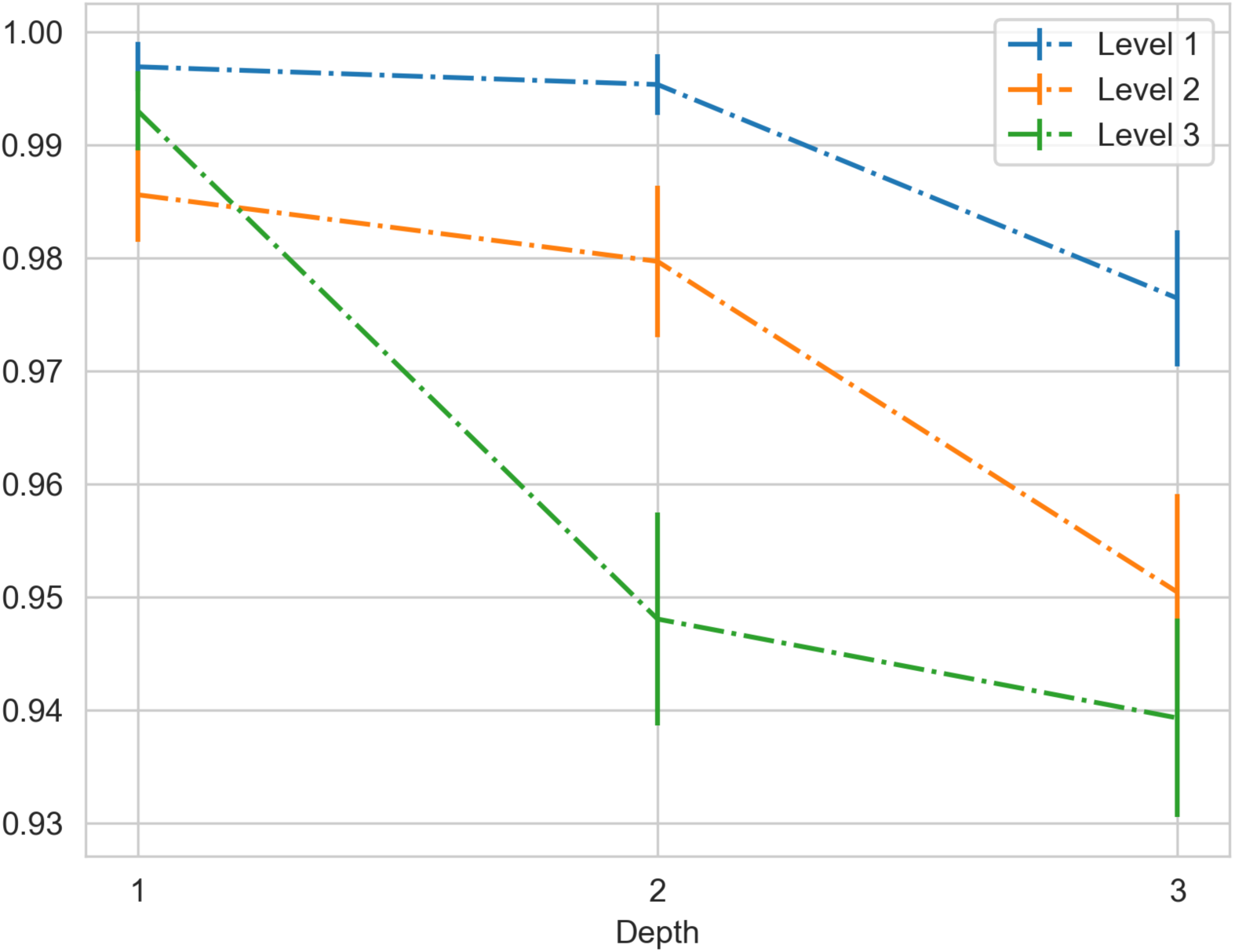
Success probability as a function of the 3 levels of the experiment and of the 3 depthIDs. The figure permits appreciating that participants’ performance decereases as a function of levels, but remains stable across levels when the problem planning depth is 1. See the main text for details. Error bars have been assigned according to the standard error of the mean of a binomial distribution.

**Figure 3.**
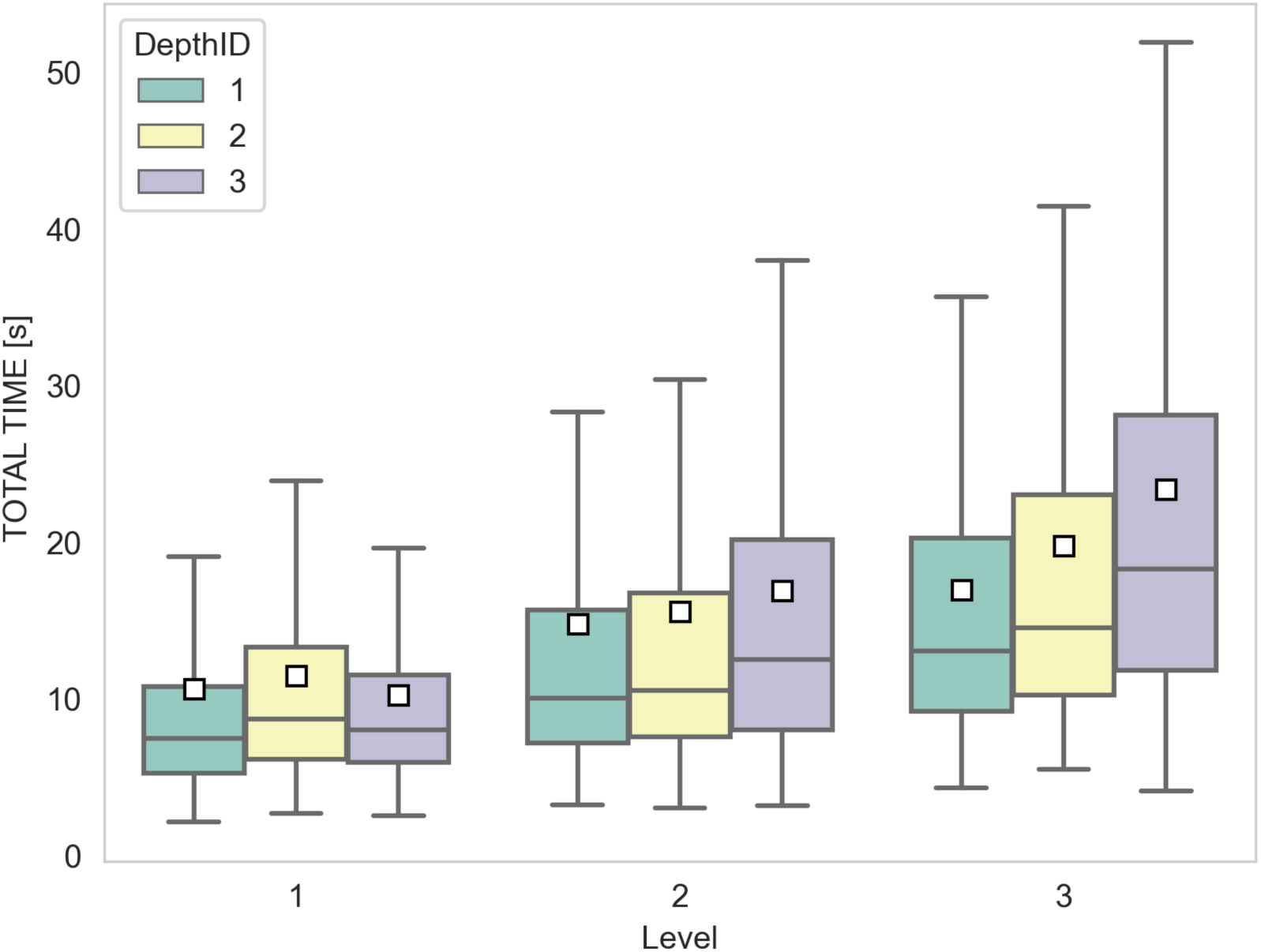
Problem completion time (in seconds) as a function of the 3 levels of the experiment and of the 3 planning depths. Results are organized by level and depth. The plot shows the total time, which is bounded to 60 seconds in our experiment.

**Figure 4.**
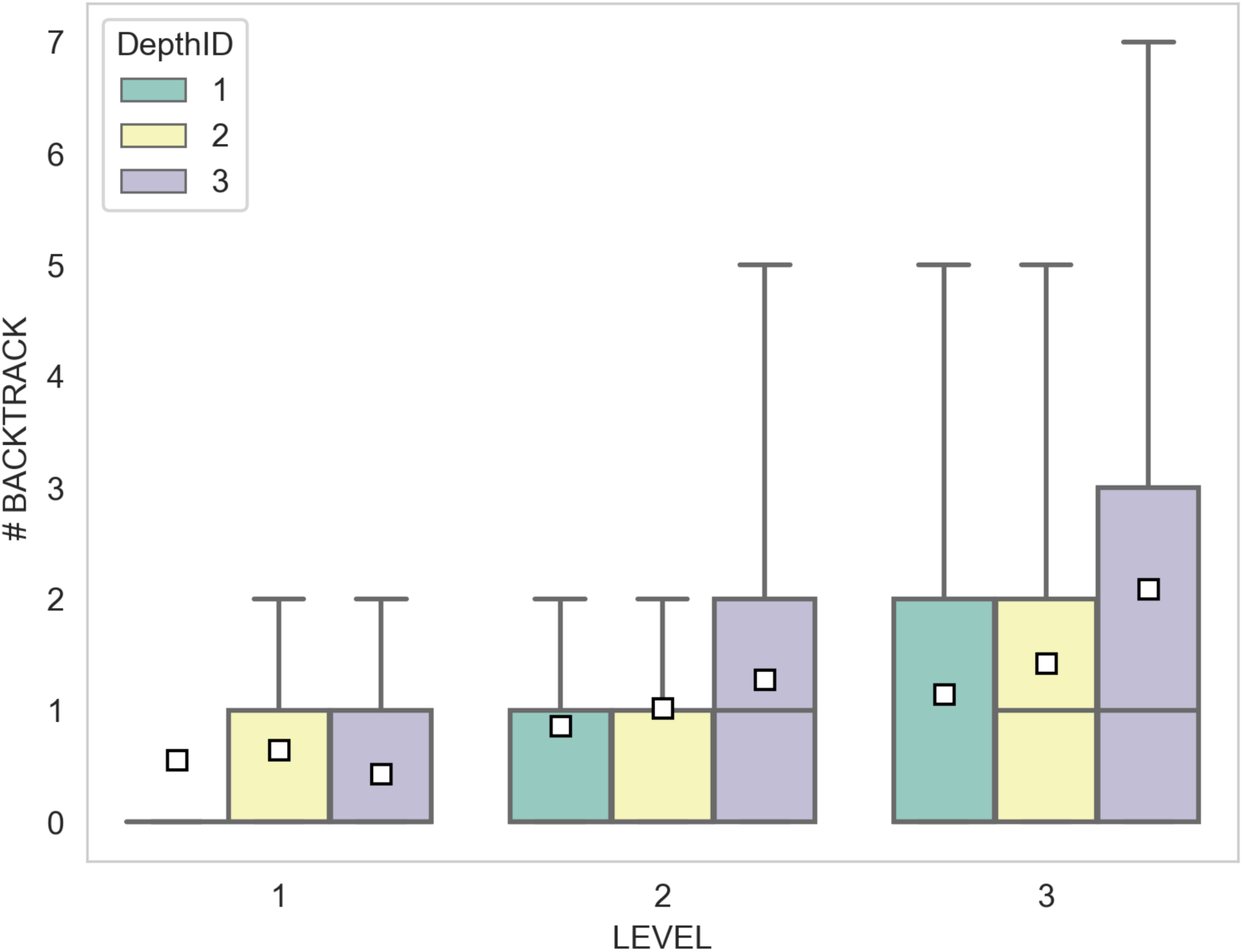
Total number of backtracks as a function of the 3 levels of the experiment and of the 3 planning depths. Results are organized by level and depth.

Crucially, we designed problems that require different planning depths to be solved. The concept of planning depth has not a unique definition but is in general used to refer to the property of “how much further from my current state am I thinking of”. In this work, we anchor the planning depth to the number of gems (from 1 to 8) that a person considers collecting in the current plan. Some problems could be solved using a “greedy” strategy to always move to the closest gem (i.e., planning depth 1). However, in other problems, following the “greedy” strategy leads to a dead end. To solve these problems, it is necessary to select a plan to move to the next 2 to 8 closest gems (i.e., planning depths 2 to 8). For example, the problem shown in Figure 1 requires a planning depth of 5. This means that finding a(ny) solution requires planning 5 gems in advance and any planner that only considers a smaller number of gems would fail. Henceforth, a problem is said to require planning depth *n* if *n* is the minimum number of closest gems that an agent has to consider in the plan, in order to be able to solve the problem.

This design allows us to study the planning depth that participants use to solve each problem, by systematically comparing their behaviour with the behaviour of 8 variants of the same planner, which only differ for the depth *n* parameter (with depth *n* ranging from 1 to 8). In our analysis, we focused solely on the paths chosen by participants before they made their first backtracks, as these paths serve as valuable indicators of their initial plans. This approach also allows for easier alignment with findings from studies on other games like chess, where movements cannot be reversed.

If people always use the same planning depth (e.g., depth 3) to solve all the problems, then their behaviour should always match the same planning model (e.g., the one at depth 3) across all problems. If instead people recruit resources according to task demands, they will adapt their planning depth to the minimum depth required to solve the problem, and then their behaviour should match a different planner for each set of problems; namely, the planner that uses the (set-specific) minimum depth.

To preview our results, we found that people tend to use the minimum depth required to find the solution to any specific problem, indicating that they flexibly adapt their planning resources to the situation.

## Methods

### Data Collection

The experiment was conducted with the support of ThinkAhead, an Android application developed to study navigational planning and problem solving (https://play.google.com/store/apps/details?id=com.GraphGame.Conan). We recruited 160 participants online and all gave informed consent to our procedures which were approved by the Ethical Committee of the National Research Council (Commissione per l’Etica e l’Integrità nella Ricerca, Ethical Clearance protocol n. 0072130/2019, 18/10/2019). Participants were free to leave the experiment at any moment. In the analysis, we consider the 65 participants (42 male, age = 34 +- 11 years; 19 female, age = 35 +- 10; 4 participants who preferred not to specify their gender, age = 34 +- 10 years) who tried at least 80 of the 90 problems of the experiment, although they did not necessarily solve all of them.

### Experiment design

The experiment comprised 90 problems, each requiring participants to collect all the “gems” (i.e., colored dots) in a grid, without passing twice through the same node, in 60 seconds; see Figure 1 for an example problem and Figure S1 for other example problems of the three levels and planning depths. Participants were instructed that they would earn points proportional to the time left to solve the problems and that the points were doubled in problems where the gems were red (which happened in half of the trials) compared to those where the gems were blue (in the other half of the trials). As soon as the problem was shown to the participants, the time countdown started. If the participants did not solve a problem within the deadline, it counted as a failure; participants were allowed to either complete it (without getting any points) or to skip it and pass to the next one.

We generated a range of problems that required different planning depths, from 1 to 8 (see the next section for the exact definition of planning depth). The problem grids were all unique and were generated with an average level of edges density of 0.75 +- 0.2 (i.e., on average, 25% of all the possible edges where removed), which we found in a pilot study to afford a good range of planning solutions.

Before the experiment, participants performed a short practice session, in which they had to solve 4 problems, whose results were not analysed. The 90 problems were divided into 3 interleaved blocks (henceforth, “levels”), with 30 problems for each level. We varied planning demands both within and between levels; see Table 1 and Supplementary Figure S1. To vary planning demands between levels, the 3 levels were characterized by increasingly large maps and more gems to be collected, making the higher-level problems (on average) more challenging. To vary planning demands within levels, we divided each level into 3 sublevels of 10 problems each. The 3 sublevels comprised problems that could be solved using planning depth 1, 2-4, or 5-8, respectively; see Table 1. Note that the levels have (on average) increasing difficulty but include a fixed proportion of easier trials (i.e., requiring lower planning depth). This design permits increasing the challenge and simultaneously maintaining a high frequency of success across levels, which is important for intrinsic motivation (52).

**Table 1.**
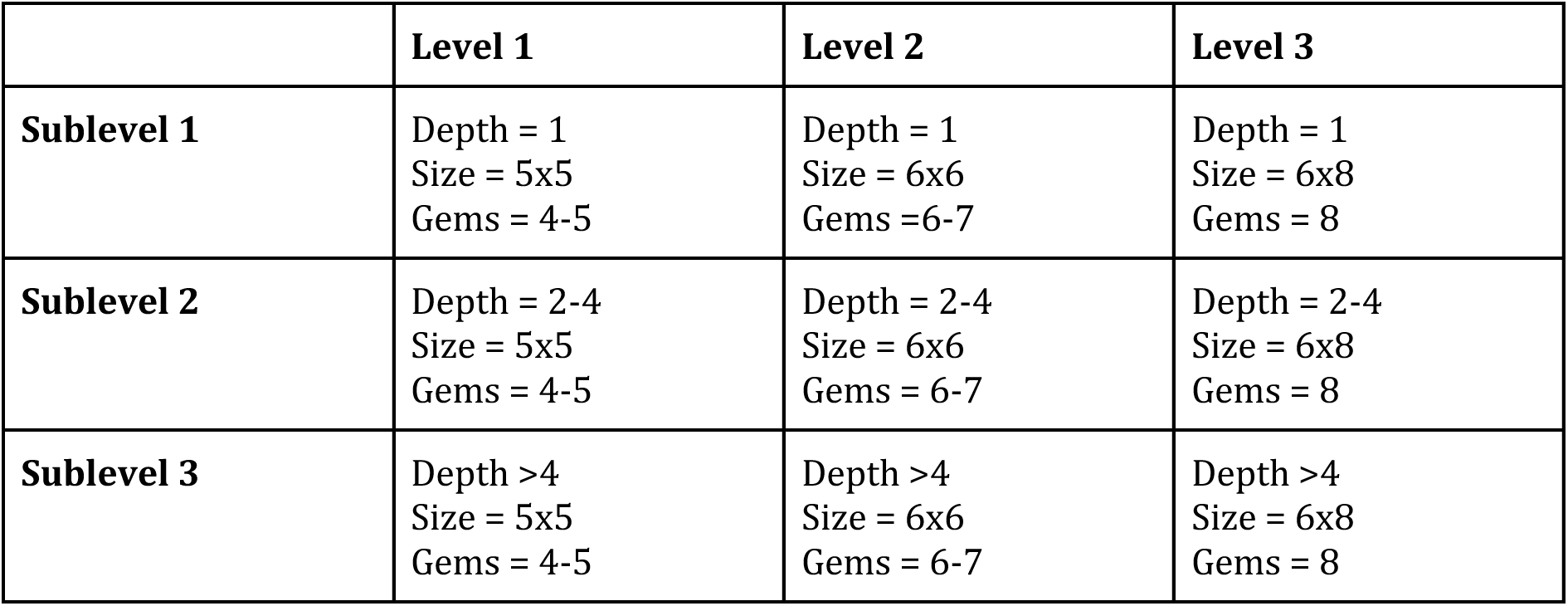
Experiment design. The experiment is structured into 3 levels of increasing (average) difficulty, each including 30 problems. See the main text for explanation.

### Assessment of the minimum planning depth required for each problem

To assess participants’ planning depth during the experiment, we designed a planner that computes all the simple paths (i.e., paths that do not pass twice on the same node) that start from the current position and reaches *n* gems. In our analyses, we consider 8 variants of the same planner, which only differ for the *n* parameter, which represents planning depth (from 1 to 8) (17,18). For example, a “greedy” planner with *n* = 1 will look for all the simple paths that from the current node reach one and only one gem (a planner with *n* = 2 will look for all the simple paths crossing only 2 gems, and so on). After the computation of all these possible (partial) paths, the shortest one is selected. If there are many equivalent shortest paths the choice will be uniformly random among them. The last position of the selected path becomes the new current position, and all the crossed from the chosen paths are tagged as *visited*, and thus removed from the computation of future possible paths to comply with task rules. After this, the computation of the next (partial) paths is repeated, until all the gems have been collected, or there are not enough gems that the agent is able to collect (i.e., less than the minimum between its planning depth and the remaining gems). The planners considered so far cannot backtrack, so once a dead-end is reached, the simulated trial ends. Note that since there might be various paths to choose among, multiple runs of the same planner on the same problem might have different outputs; for example, because only one of the shortest paths selected at a certain point will lead to the solution of the trial. See Algorithm1 in the Supplementary Materials for the pseudocode of the planning algorithm and the routine “AgentForward.py” in the Data and Code repository for the detailed script.

We used the 8 variants of the planner to assess the minimum planning depth required to solve each problem. For this, we classified each problem according to the minimum value of *n* for which a solution could be found with non-null probability, from 1 to 8. For example, problems of depth 5 can only be solved by a planner with *n* = 5 (or possibly more), but not by a planner with *n* = 1 to 4.

This permits us to group the problems into 8 groups, with the index denoting the minimum planning depth required to solve them. Note that the fact that a problem can be solved by a planner with depth *n* does not imply that deeper planners can necessarily solve the same problem, or that planners at different depths find the same solution to the same problem. Let’s suppose, for example, that a solution to a problem exists for a planning depth of 5. This solution reflects the shortest path connecting 5 gems, but there is no guarantee that a planner at depth 6 (or higher), which looks for the shortest way of connecting 6 gems, would find the same solution. This might be the case, for example, when a longer path to the first 5 gems is needed in order to reach the sixth one with a globally shorter solution. Notable, the fact that planners having different depths could solve the same problem in different ways permitted us to estimate participants’ depths more carefully (see Supplementary Figures S5 - S12).

### Analysis of success probability, problem completion time and total number of backtracks

We used multiple logistic and linear regressions and the R library lme4 (53) to analyse participants’ success probability (i.e., the probability that they solved the problems before the deadline of 60 seconds), problem completion time (i.e., the average time they needed to complete the problem, in seconds) and total number of backtracks that they executed during the experiment. The 3×3 design considers the 3 levels of the experiment (Levels 1-3) and the 3 planning depths (DepthID 1-3, corresponding to depth 1, depths 2-4, and depths 5-8, respectively).

### Similarity between participants and planning models, for each of the 8 problem groups

The comparison between human participants and planning models involved assessing the similarity in planning depth across the eight problem groups that required minimum planning depths ranging from 1 to 8. For each of the 90 problems, we determined the likelihood of the data (number of gems collected by the participants before the first backtrack) given the models (the eight planning models). To ensure comprehensive coverage of potential solutions at varying depths, we utilized 500 instances of each planner for every problem. Subsequently, we identified which planner, with a planning depth ranging from 1 to 8, exhibited the highest probability (i.e., was the ‘winner’) within each problem group.

## Results

### Participants are sensitive to both problem level and planning depth, but in different ways

We assessed the statistical significance of the effects of the experimental design (Level x DepthID) on the success probability (i.e., the probability that they solved the problems before the deadline of 60 seconds). For this, we used a general linear mixed-effect model with Level and DepthID and their interaction as fixed-effects while accounting for subject identity variability as a random effect. The values of the coefficients of the model are reported in Table 2, with their errors and significance. We were able to assess the statistical significance of the different terms of the full model by comparing it against reduced models that excluded some fixed effect. This shows significance for the main effect of the DepthID and the Level terms, while no interaction was found. Also, the significance of the use of the random term was assessed comparing the full model with one without the random term: (χ^2^(1, 5697) = 104 p- value < 2.1*10^-16^).

**Table 2.**
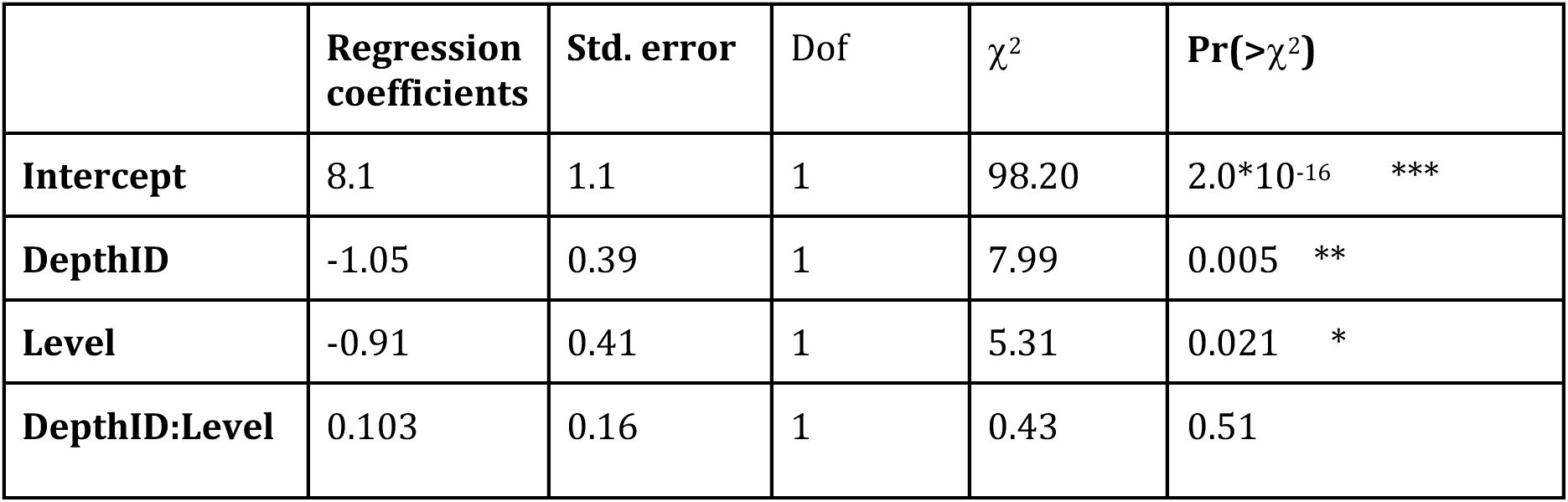
Success probability regression. A general linear mixed-effect model is used to fit success probability. Significance code (0< *** <0.001; 0.001** <0.01; 0.01 < * <0.05).

| | **Regression coefficients** | **Std. error** | **Dof** | $\chi^2$ | **Pr(>$\chi^2$)** |
| --- | --- | --- | --- | --- | --- |
| **Intercept** | 8.1 | 1.1 | 1 | 98.20 | 2.0*10<sup>-16</sup> *** |
| **DepthID** | -1.05 | 0.39 | 1 | 7.99 | 0.005 ** |
| **Level** | -0.91 | 0.41 | 1 | 5.31 | 0.021 * |
| **DepthID:Level** | 0.103 | 0.16 | 1 | 0.43 | 0.51 |

We also used a linear mixed-effect model to fit the relation between the completion time and the experimental design, with DepthID, Level and interaction as fixed effects and the subject identity as random effect. The coefficients of the regression are shown in Table 3, with their standard errors and their significance. The inclusion of the random effect was shown significant by a model comparison of the full model with and without it ( χ^2^(1,5697) = 853, p-value < 2.1*10^-16^). In an analogous way, we assessed the significance of the terms by a comparison of the full model with reduced versions.

**Table 3.** Completion time regression. A linear mixed-effect model is used to fit completion time. Significance code (0< *** <0.001; 0.001** <0.01; 0.01 < * <0.05).

| | Regression coefficients | Std. error | DoF | $\chi^2$ | Pr(>$\chi^2$) |
| --- | --- | --- | --- | --- | --- |
| **Intercept** | 4.7 | 1.3 | 1 | 12.43 | 4.2*10<sup>-4</sup> *** |
| **DepthID** | 0.93 | 0.52 | 1 | 3.18 | 0.075 |
| **Level** | 2.76 | 0.53 | 1 | 26.93 | 2.1*20<sup>-7</sup> *** |
| **DepthID:Level** | 0.91 | 0.24 | 1 | 14.16 | 1.7*10<sup>-4</sup> *** |

A linear mixed-effect model for the number of backtracks (significance of the addition of the random effect: ( χ^2^(1, 5697) = 274, p-value < 2.2*10^-16^) shows no main effect of DepthId and Level, but a significance for the interaction term, see Table 4.

**Table 4.** Number of backtrack regression. A linear mixed-effect model is used to fit number of backtracks. Significance code (0< *** <0.001; 0.001** <0.01; 0.01 < * <0.05).

| | Regression coefficients | Std. error | DoF | $\chi^2$ | Pr(>$\chi^2$) |
| --- | --- | --- | --- | --- | --- |
| **Intercept** | 0.046 | 0.18 | 1 | 0.066 | 0.80 |
| **DepthID** | 0.011 | 0.077 | 1 | 0.0198 | 0.88 |
| **Level** | 0.14 | 0.078 | 1 | 3.13 | 0.077 |
| **DepthID:Level** | 0.18 | 0.035 | 1 | 25.95 | 3.5*10<sup>-7</sup> *** |

Finally, a linear mixed-effect model for the view time shows no significant main effect of DepthId or Level and no significant interaction (Figure S16 and Table S1).

### Participants’ initial planning depth is adaptive and matches task demands

We assessed the similarity between the initial plans (i.e., the plans before the first backtrack) of participants and planning models, for each of the 8 problem groups. The results of this analysis, aggregated across the three levels and for red and blue gems, are shown in Figure 5. Our results indicate that the planner that best explains the behaviour of participants (i.e., has the maximum likelihood for the majority of problems) is the one having the minimum required planning depth. For example, the planner that best explains the behaviour of participants during the solution of problems requiring minimum planning depth of 1 is the planner using depth 1. The same similarity between minimum planning depth of the problem and of the planner that best explains participants’ data is observed across all the 8 levels, see Figure 5.

**Figure 5.**
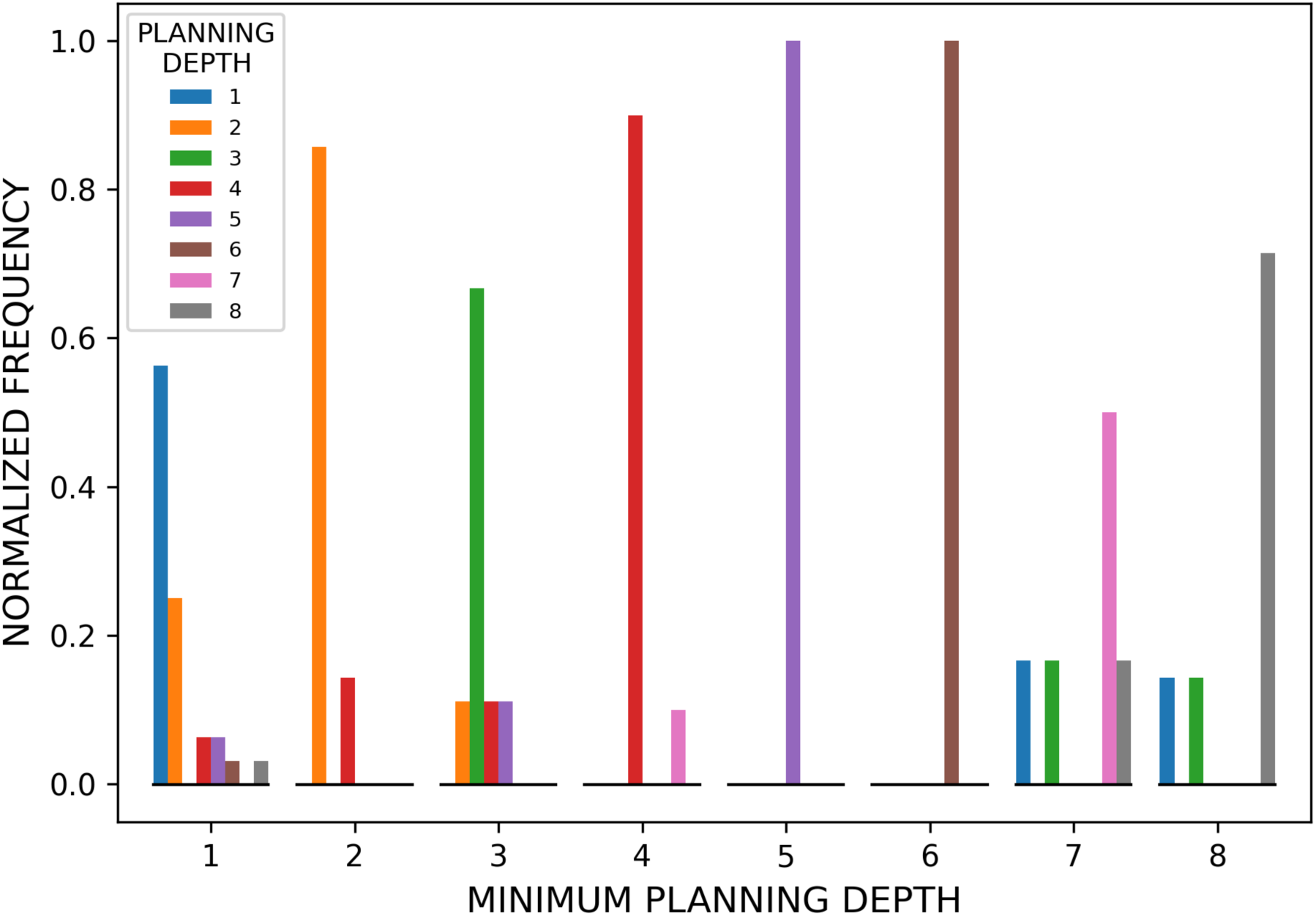
Comparison between participants and planning models. The figure shows the number of times that each planner from depth 1 to 8 had the maximum likelihood of the gems collected before the first backtrack by the participants, in the problems of each of the 8 problem groups. Problems are grouped according to the minimum planning depth required to solve them, from 1 to 8, and color coded (see the legend). The frequency of each planner is normalized within each group. The figure shows that for each set of problems, from 1 to 8, the planner that wins most frequently (i.e., that has the greatest likelihood in the plurality of simulations) is the one having the minimum planning depth.

In order to assess the statistical significance of this consideration, we tested how well using a fixed planning depth described participants’ behaviour against a model where the planning depth was adapted to the minimum planning depth required by the problem. To do that, we computed for each problem the distribution of the rankings in the likelihood obtained by each possible value of the planning depth (e.g., if in a problem the model using a planning depth of 2 has the highest likelihood, it has a rank of 0 for that problem; all the other planners are ranked in a decreasing order, accordingly to the same procedure). So, we obtained a distribution of the ranks for each planning depth (90 values each). We also computed the rank of a planner that adapts its planning depth to the minimum required for that problem. The resulting distributions are shown in Figure 6.

**Figure 6.**
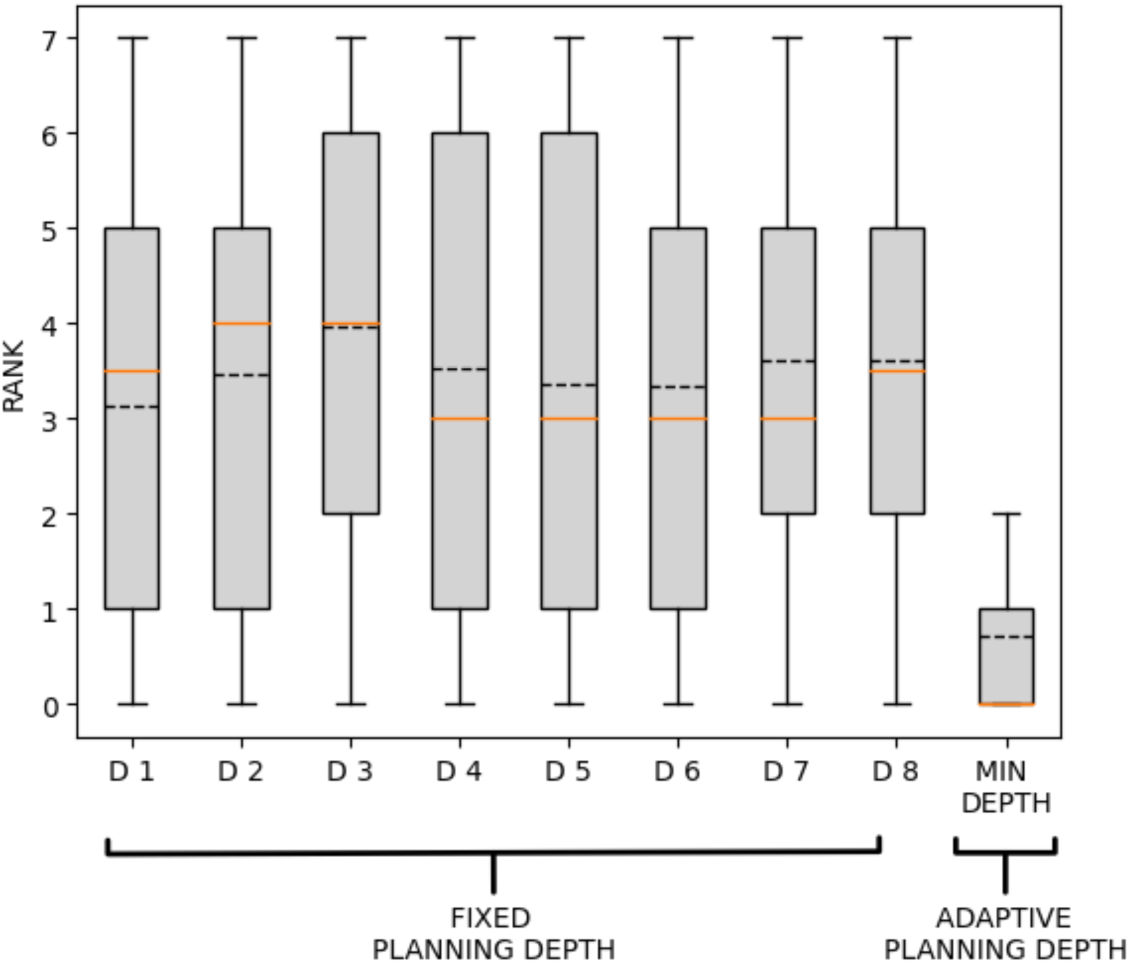
Rank values of fixed and adaptive depth planners across the 90 problems. The figure shows the distributions of the ranks of planners using a fixed planning depth (D1-D8) across all problems and of the planner using a planning depth adapted to the minimum required depth to solve a problem (Min depth). The black dotted lines show mean values, whereas the orange lines show median values. See the main text for explanation.

To test whether the average rank of the adaptive planning depth model (min depth in Figure 6) is significantly lower than each of the fixed planning depth models (D1 to D8 in Figure 6), we performed chi squared independence tests. We found that for each couple (min depth – D1, min depth – D2, etc.), the rank distributions are significantly different (χ^2^(1, 90) = 57, p-value <4.0*10^-10^). This implies that adaptive planning depth to the minimum requirements of the problem provides the most accurate description of participants’ behavior. To rule out the possibility that the result could be dependent by the possibility of ties between different depths on the same problem, we performed 10^5^ iterations of the same test, in which we added a small Gaussian noise to the likelihoods of data according to each planning depth. We found that our claim was rejected only 0.0065% of the times, therefore indicating the robustness of our results.

An equivalent description to the log-likelihood computation (though visually more compact) is obtained when considering the Kullback–Leibler (KL) divergence between the distribution of the number of gems collected by the whole group of participants for any given problem before the first backtrack, and the distributions of the number of gems collected by 500 instances of each of the 8 planners (see Supplementary Figure S2). We next tested for the robustness of these analyses and obtained the same trend when examining the problems separately for the 3 levels of the experiment (Supplementary Figure S3), separately for red or blue gems (Supplementary Figure S4) and grouping them into classes reflecting their minimum required depth and maximum number of gems (Supplementary Figures S5 – S12). These control analyses indicate that our main findings do not depend on map size (which varies across levels and class groups) or incentives (which are different for red and blue gems). Furthermore, these control analyses highlight that for all problem groups, the 8 planners collect different average gem distributions and are therefore distinguishable (Supplementary Figures S5 – S12).

To test the robustness of the above results, we asked whether they remain the same when using other types of planners with varying planning depths. First, we considered an “obvious ending” planner: a variant of the planner used in the main analysis that knows the identity of the last gem to be collected. The motivation the “obvious ending” planner comes from the fact that some of the problems in our experiment (n = 44) had an obvious ending: if a gem was placed in a dead-end or close to it, it was necessarily the last one to be collected. We incorporated this information in the “obvious ending” planner, by calculating all the paths as in the planner used in the main analysis but excluding all the paths where a gem in a dead end (or was only connected to dead ends) was not the last one in the plan. This procedure leads to 8 variants of the “obvious ending” planner, one for each depth level. We then compared the performance of the 8 planners used in the main analysis and of the 8 “obvious ending” planners, by computing for all the problems how many times each model was the best fitting of participants’ performance, across all problems. We found that in the problems where the two model where distinguishable, the best description was most frequently provided by the planner used in the main analysis, even though the frequency difference was minimal, see Figure S13. Finally, we found that the advantage of using an adaptive planning depth over a fixed planning depth emerges also when considering the “obvious ending” planners, showing that it is robust to information about the last gem to be collected (see Figures S14 and S15).

We next considered planners that introduce noise or stochasticity in action selection. The motivation for this choice is the observation that the simple, almost deterministic planners used in the main analysis fit well the most probable number of gems collected by participants at each planning depth, but not the entire distributions of collected gems (Figures S5 – S12). This suggests a source of stochasticity or noise in participants’ choices that is not well captured by the (mostly) deterministic planners used in the main analysis. To understand which source of noise best explains participants’ behaviour, we compared two extensions of the planner used in the main analysis, which introduce noise at two different stages of the planning problem. The first extension is an ϵ-greedy planner that selects the next node randomly (rather than using planning) with a certain probability ϵ, which is a free parameter. The second extension is a Softmax planner that replaces the Argmax action selection used in the main analysis with a Softmax function, whose temperature β is a free parameter (see the Supplementary Materials for pseudocode of the two planners). Note that the planned used in the main analysis is a special case of the two planners, having no noise ϵ and a very high inverse temperature β. We compared the two planners using maximum likelihood estimation and found that the ϵ-greedy planner outperforms the Softmax planner in fitting participants’ data in all the problems. Furthermore, we found a significant linear relation (p_pearson_ = 0.6, p-value < 10˄-9) between the optimal ϵ of the ϵ-greedy planner and the planning depths of the problems (Figure S17). This result indicates that the increase of planning depth of problems is associated with an increase in the optimal noise required to fit the data – and that participants’ behaviour appears more stochastic and possibly more suboptimal when facing problems requiring greater planning depth. Finally, we tested whether the noisy (ϵ-greedy) planner showed the same adaptive planning depth of the main analysis. We found that, analogous to the main analysis (Figure 5), the ϵ-greedy planner that best explains the behaviour of participants (i.e., has the maximum likelihood for the majority of problems) is the one having the minimum required planning depth (Figure S18). Furthermore, analogous to the main analysis (Figure 6), the average rank of the adaptive ϵ-greedy planning depth model is significantly lower than each of the fixed ϵ-greedy planning depth models, except for the fact that it is indistinguishable from the planner using depth D8 (Figure S19). These results largely replicate the adaptive planning depth of the main analysis, even in the case of noisy (ϵ-greedy) planners.

### Participant prefer occupy nodes well connected to other well-connected nodes

So far, we focused on problem solving at the level of selection between goals and subgoals (i.e., gems). However, the same subgoal can usually be reached through multiple paths. To get more insight on the paths selected by participants, we considered their node occupancy during the solution of the problems. We removed from the analysis all the forced choices, i.e., the nodes that were part of any possible solution, such as the start positions and the gems. We next calculated seven graph-theoretic measures for the nodes, averaging across all problems (54). These include 1) *eigenvector centrality*, representing the influence of a node in a connecting network, which has a high score if a node is connected to many nodes having themselves high scores; 2) *relevant goal information*, defined as the information needed in a certain node to reach the next gem (55,56); 3) *closeness centrality*, calculated as the reciprocal of the sum of the length of the shortest paths between the node and all other nodes in the graph; 4) *degree centrality*, defined as the number of edges of the node; 5) *random*, which assigns the same likelihood to all nodes belonging to a possible solution; 6) *betweenness centrality*, defined by considering the number of shortest paths to any other node that pass through a given node; and 7) *shortest path*, which assigns a higher value to nodes that are more often part of a shortest solution. We converted each metric into a probability distribution of node occupancy, using a Softmax of the scores of the metric; for example, the model of the occupancy based upon the eigenvector centrality assigns a greater probability to nodes with higher values of the eigenvector centrality metric. We compared the log likelihood of participants’ occupancy of the nodes according to the seven graph-theoretical metrics and we found that the best fitting graph-theoretic measure (having the highest log likelihood) is eigenvector centrality, whereas the worst fitting measure is shortest path. The results of the model comparison are shown in Figure 7, as the negative log likelihood excess between the best metric, eigenvector centrality (having zero excess), and all the other metrics.

**Figure 7.**
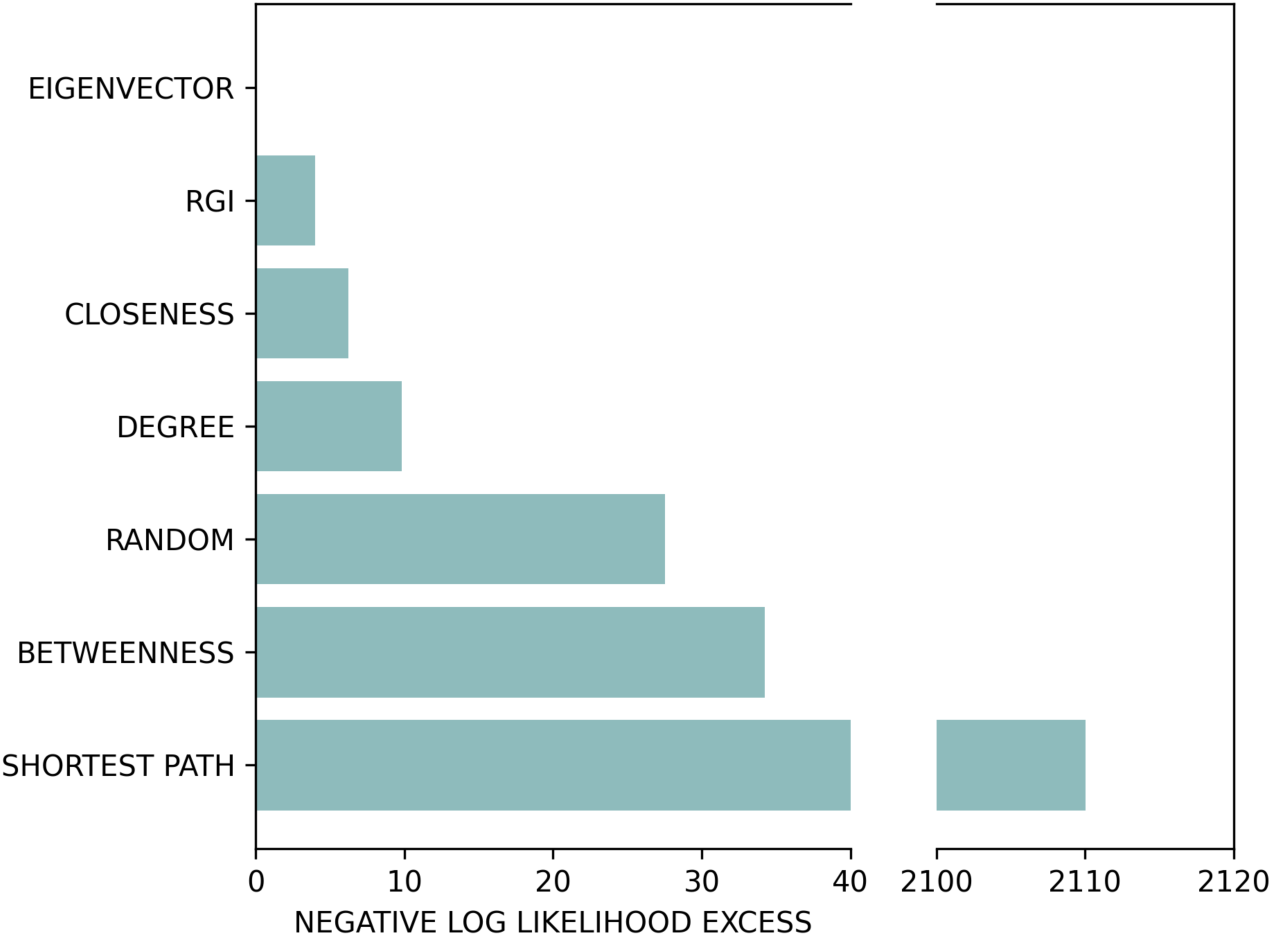
Log likelihood of participants’ node occupancy during the solution of the problems, using seven graph-theoretical metrics. The metrics are eigenvector centrality (EIGENVECTOR), relevant goal information (RGI), closeness centrality (CLOSENESS), degree centrality (DEGREE), random node occupancy (RANDOM), betweenness centrality (BETWEENNESS), nodes belonging to the shortest path only (SHORTEST PATH). The figure shows that the best fitting metric (having the lowest negative log likelihood) is EIGENVECTOR, whereas the worst fitting model (having the greatest excess) is SHORTEST PATH.

## Discussion

Since the early days of cognitive science, researchers have asked how we solve complex planning problems that defy the exhaustive evaluation of all possible choices (1). It is commonly assumed that planning is a form of cognitive tree search using a mental map (4–8). However, except in the simplest cases, exhaustive search is infeasible, hence pointing to *bounded* forms of planning that adapt cognitive resources to task demands (13–15).

In this study, we asked whether participants adapt their planning resources to the demands posed by problems of different complexity. To address this question, we asked participants to solve a series of planning problems that required finding a path to connect all the “gems” in a grid, without passing through the same node twice. The problems were divided into three levels, characterized by increased map size and number of gems. We varied planning demands both within and between levels, therefore designing 8 groups of problems that required a minimum planning depth from 1 to 8. The planning depth required for each problem was unknown to the participants.

Our results indicate that participants’ problem-solving performance depends both on planning depth and the level of the problems, underscoring that the challenge lies both in the cognitive demands imposed by a problem and its spatial scale (which increases with problem level), independently. Furthermore, the time required for problem-solving is affected by spatial scale, illustrating that addressing problems within larger maps inherently demands more time. Also, the effect of the planning depth on time required for problem-solving becomes relevant, via the interaction with the spatial scale. One possible explanation is that while forming a complete plan early on is possible with smaller maps and simpler problems, working memory or attention limitations might preclude this with larger maps and more challenging problems. In other words, simpler and smaller problems would afford a global evaluation strategy, while more challenging and larger problems would require instead a more local evaluation strategy. Additionally, the analysis of the number of backtracks shows that the spatial scale of the map and the complexity of the problem increase how often participants need to change their initial plans, but the effect size is not independent for the two dimensions.

Crucially, our results show that participants flexibly adapt their initial planning depth (i.e., the depth of their plans before the first backtrack) to the minimum depth required to solve the problems, as estimated using planning models (Figure 5). The adaptiveness of participant’s planning depth is confirmed by the fact that a planner that uses the minimum depth level (from 1 to 8) for each type of problem better explains participants’ behaviour, compared to a(ny) planner that uses a fixed planning depth across all problems (Figure 6). This result is robust to various control analyses: it is observed across all the 3 levels of the game (Figure S3) and the 2 types of gems, blue or red, associated to lower and higher rewards, respectively (Figure S4), when considering planners that know the identity of the last gem to be collected, when this information is obvious to participants (Figures S13-S15) and (ϵ − *greedy*) planners that perform noisier action selection (Figures S18-S19). Finally, our results indicate that in the selection of paths to reach the next planned gems, participants follow prefer occupying nodes that are well connected to other well-connected nodes (i.e., have high eigenvector centrality), even when these are not part of the shortest path. This result might indicate a possible strategy or heuristic to avoid getting stuck in dead ends.

Taken together, these results suggest that during problem solving, people make an adaptive use of their cognitive resources, by selecting an appropriate level of planning depth. Therefore, this study adds to a large literature – starting with the pioneering work of Simon – showing that during problem solving, people adapt to the complexity and structure of the environment (24,27,57) and make an adaptive use of their cognitive resources (58). Furthermore, the fact that participants prefer occupying well connected nodes rather than nodes belonging to the shortest path is in keeping with studies showing the adoption of approximations to optimal solutions and heuristics, especially when the problems are challenging (24,25,59). For example, people prune unpromising branches of the search tree (19,20,60) and reduce tree search under time pressure (21). An emerging idea is that these (and other) approximations to optimal solutions might stem from a “rational” and efficient use of limited resources– i.e., *bounded* or *resource-rational* planning (46,47). For example, during problem solving, people might spend more time and effort planning ahead when the benefits of investing cognitive resources are greater and when the problem requires more deliberation (61–63).

An open objective for future research is assessing to what extent the results reported in this study generalize to other planning tasks. An advantage of our task is that it incentivizes planning, as opposed to (for example) moving as fast as possible to find a solution by change, because the problem space is relatively large, the subgoals (gems) are clear, there is a time incentive, and backtracking adds a time penalty. The combination of these factors might have encouraged people to plan ahead in adaptive ways, as testified by the good match with (deep) planners. At the same time, our task is not representative of all the conditions in which planning can arise, because it is static and (at least in principle) fully observable. Future studies might address situations in which certain task elements, such as edge positions or gem values, change dynamically over time, possibly requiring “detours” (64). Such dynamic environments require considering how to adapt action plans to varying temporal demands and how to replan when an existing plan becomes unattainable. Similarly, future studies might address situations in which certain task elements, such as the presence or absence of a particular edge, are initially unknown or known only with a certain probability, as in the Canadian Traveller Problem (65). Such partial observability requires balancing exploration and exploitation during planning (66,67). Future studies might assess whether the results reported in this study generalize to dynamical and partially observable settings, or to settings that present participants with different incentives for planning versus following habits (56).

This study has various other limitations that need to be addressed in future work. First, this study indicates that participants adapt their initial planning depth to task demands but does not clarify how they do that. There are multiple alternative strategies that could explain our findings. For example, before navigation begins, participants might use a “gist” of the maze to decide planning depth – or how much cognitive resources they would need to invest to plan ahead – and then use such planning depth to find a solution (note that selecting an appropriate planning depth does not necessarily entail solving a particular problem, because there are usually several alternative paths at the same planning depth). In the planning literature, there is a key distinction between an “encoding” phase in which participants form a mental representation of the problem and the “planning” phase, in which they form a plan based on the mental representation. Previous results suggest that during the encoding phase, participants might form simplified mental representations of the problems, which omit task-irrelevant details rather than being complete (68). It is possible that this simplified (or gist) representation could be sufficient to guide the selection of an appropriate planning depth, but future studies are required to understand whether and how this is possible. Alternatively, before navigation begins, participants might start searching for a solution at low planning depth and then increase the depth progressively, until they find a (satisficing) solution. Future studies looking at eye movements before and during problem solving could contribute to address these questions (69).

Another limitation is that this study does not address the “algorithm” used by the brain to solve the problems. When establishing a similarity between the planning models and human participants, we are not necessarily claiming that human participants use the simple planning algorithm described in the paper, but only that they appear to adapt their planning depth across problems – regardless of mechanism (e.g., a different planner for each problem, “iterative deepening depth-first search” (39,70) that start from an initial “cheap” plan and progressively refine it (56,71,72), or other methods). Having said this, our specific setting, in which the problem maps are novel and fully visible, epitomizes the use of model-based planners (5,73–76). While in principle the problems considered in this experiment could be solved using model-free and successor representation algorithms that dispense from planning (67,77,78), these are unlikely candidates, since they would require an extensive learning phase, whereas in our experiment the participants never see the same map twice. Future studies might compare more directly various additional planning (or even non-planning) algorithms. Apart for those discussed above, another relevant class of algorithms is Monte Carlo planning. These algorithms offer an approximate solution to the problem of sampling in large or continuous state spaces (79) and have been linked to human cognitive search (33,35,62), but would behave similarly to depth-first planners in our (relatively small scale) problems. Another possibility is using hierarchical planners, which split large problems into smaller, more manageable ones (29,32,80). Given that the focus of this paper is on the first plans that people form, not on their entire plans, the use of hierarchical planners seems less compelling; but future studies could use hierarchical planners to extend the results of this study to the entire plan selected by participants. Furthermore, future studies could test the hypothesis that participants use a mixture of planning depths not only between trials, as we show here, but also within trials. Testing this hypothesis would require a different design, which includes a balanced number of trials requiring different mixtures of planning depths (rather than only different planning depths as done in this study) and in which the behaviour of planners using different mixtures of planning depths is distinguishable. Relatedly, future studies could assess the possibility that participants use different planning depths during different phases of the problem solution, such as before or after a backtracking.

This leads us to another limitation of this study: the fact that it only focuses on the initial planning phase. Focusing on the first part of the plan is meaningful, since previous research has established that the initial moves of participants during problem solving are “good enough” and revelatory of their strategy (81). Furthermore, from a methodological perspective, considering only the first part of the plan drastically simplifies the assessment of planning depth, since it does not require making assumptions about the algorithm used to decide (for example) when to backtrack or whether to change plan or plan depth along the way, as some planning models do (82). In other words, this choice does not require delving into the full complexity of planning, acting and replanning dynamics that can occur in our setup – and more broadly, within embodied or continuous decisions (83–86). However, a more complete analysis of planning dynamics in our experiments should consider that backtracking is part and parcel of human problem solving (39) and that people might generate longer plans when they are forced to do so – for example, because they cannot backtrack (87). We hope to address these open issues in future studies.

Furthermore, our experiment was not designed to study whether and to what extent participants learn novel and potentially better planning strategies over time. Future studies might address learning dynamics, by comparing how people behave when presented with different sequences of planning problems – or “training curricula” (88) – that afford or do not afford generalization across problems.

Finally, another limitation of the study is that by restricting our analysis to participants who downloaded the game and completed at least 80 problems, we could have selected those having sufficient skill and/or engagement levels. Previous studies with a much larger pool of participants reported significant individual differences in navigation ability (89), suggesting that weaker navigators might not show the same adaptive use of resources that we report here. Other, large-scale studies of navigational planning report large age-related differences (90,91). The possible differences in adaptive planning depth between good and weak navigators and across age groups remains to be tested in future studies.

## Supporting information

Video of an example level

## Acknowledgements

This research received funding from the European Union’s Horizon 2020 Framework Programme for Research and Innovation under the Specific Grant Agreements No. 945539 (Human Brain Project SGA3) and No. 952215 (TAILOR); the European Research Council under the Grant Agreement No. 820213 (ThinkAhead), the Italian National Recovery and Resilience Plan (NRRP), M4C2, funded by the European Union – NextGenerationEU (Project IR0000011, CUP B51E22000150006, “EBRAINS-Italy”; Project PE0000013, “FAIR”; Project PE0000006, “MNESYS”), and the PRIN PNRR P20224FESY. The GEFORCE Quadro RTX6000 and Titan GPU cards used for this research were donated by the NVIDIA Corporation. The funders had no role in study design, data collection and analysis, decision to publish, or preparation of the manuscript.

## Data and software availability

Data and relevant code for this research work are stored in GitHub: https://github.com/Lelumat/RS_PAPER_ThinkAhead.git and have been archived within the Zenodo repository: https://zenodo.org/records/14536992

