## Supplementary Materials for "Adaptive planning depth in human problem solving"

### Supplementary Materials: Adaptive planning depth in human problem solving

#### Supplementary figures

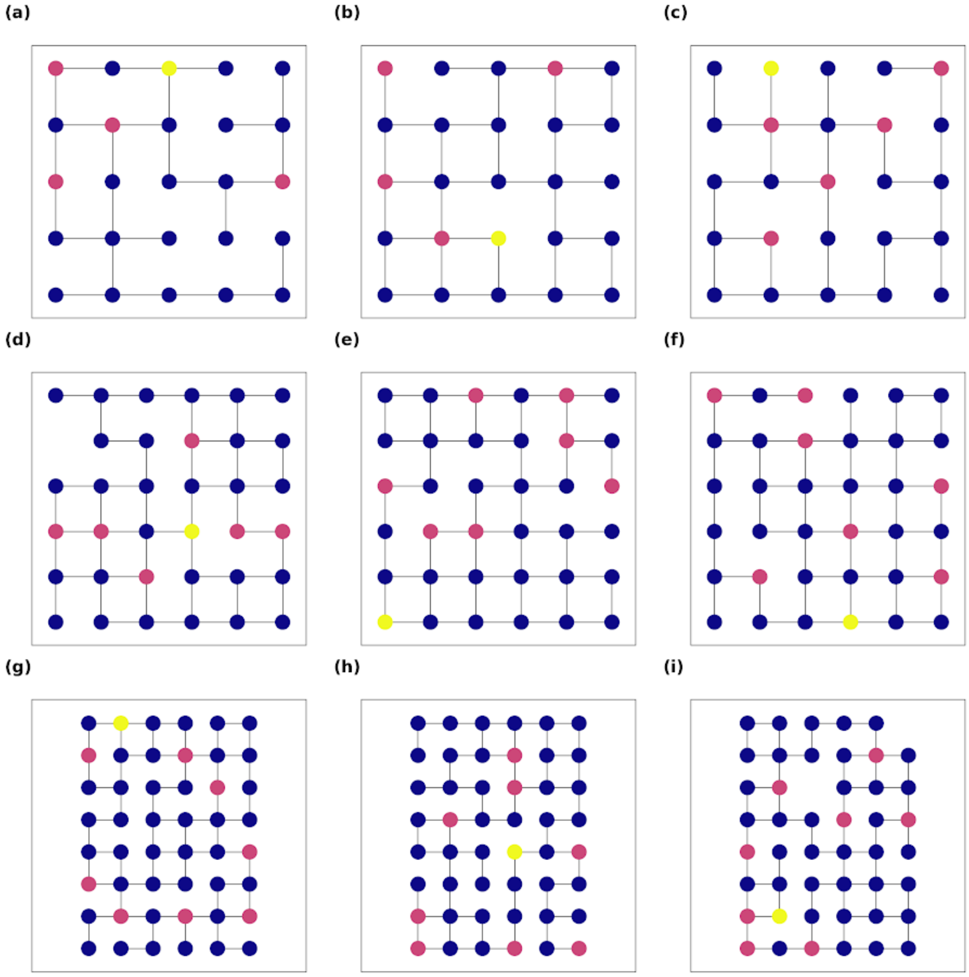


***Figure S1. Nine example problems in the experiment.*** *Each problem requires finding a path in the grid that starts from the home location (yellow node) and collects all the "gems" (red nodes), without passing twice on the same node. The 3 rows show example problems from the 3 different levels of the experiment: level 1 (a-c), level 2 (d-f) and level 3 (g-i). The 3 columns show example problems requiring different planning depths: depth 1 (a,d,g), depth 2 or 3 (b,e,h) and depth 4 or more (c,f,i). See the main text for explanation.*

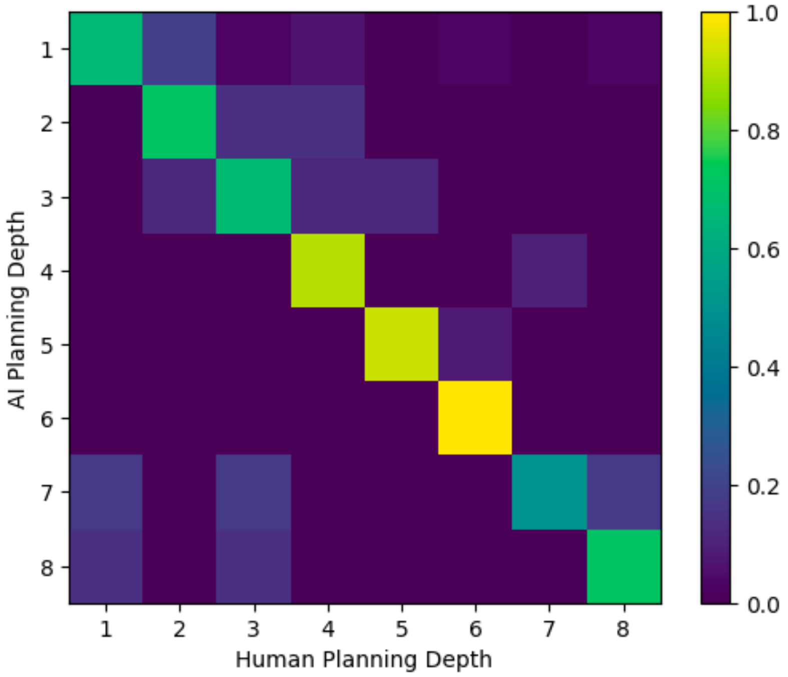


***Figure S2. Comparison of the initial planning depth of participants and planning models.*** *The row index indicates the minimum planning depth (from 1 to 8) required to solve each set of problems. The columns indicate the planning depth selected by the participants. For each row, the figure shows the probability distribution over the planning models that are most similar to the human participants, in the row-specific set of problems – with the cells having the highest probabilities indicating the best matching. Here, the metric of similarity is the Kullback–Leibler (KL) divergence between the distribution of the number of gems collected by the whole group of participants for any given problem before the first backtrack, and the distributions of the number of gems collected by 500 instances of each of the 8 planning models. The element identified by the i-th row and j-th column of the matrix represents the probability (computed as a frequency) that for a problem that required a minimum planning depth equal to i the most similar depth to humans was equal to j. For example, consider that in the fifth row, most of the probability mass is concentrated on the fifth column. This indicates that in most problems requiring planning depth 5, the behaviour of planning models with depth 5 provides the best match with the participants' behaviour. Our results show that much of the probability mass is on the diagonal or slightly above it, indicating that participants tended to select the minimum planning depth required to solve these problems, or a slightly greater depth.*

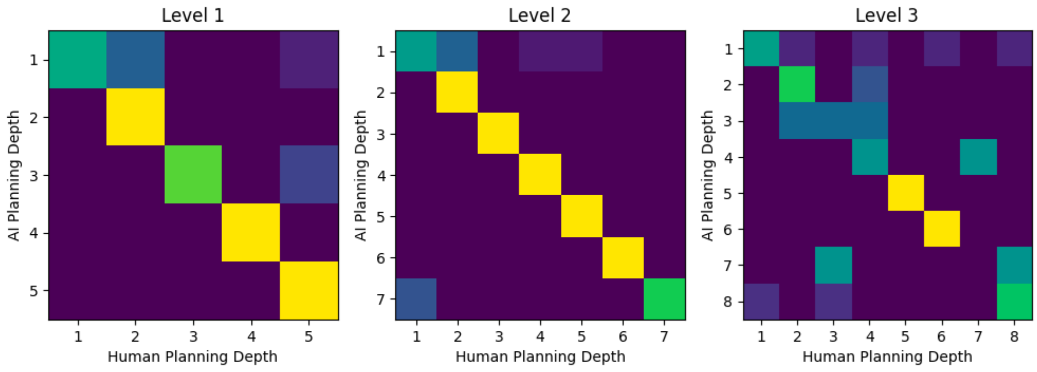


***Figure S3. Adaptive planning strategy across the 3 levels of the experiment.*** *The three figures are the same as Figure S2 but split for the 3 levels of the experiment. Please note that in levels 1, 2 and 3 the maximum planning depths are 5, 7 and 8, respectively.*

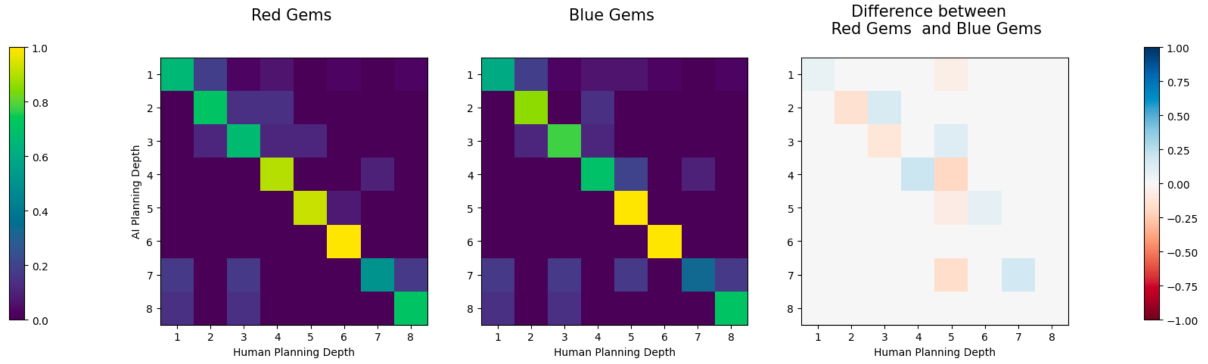


***Figure S4. Adaptive planning strategy in problems with red and blue gems, and their differences.*** *This figure is the same as Figure S2 but split for red gems (left plot) and blue gems (center plot). Finally, the differences between the first two plots are shown (right plot). While there are some differences, these do not go in one specific direction, e.g., planning depth is not consistently higher or lower for red gems across all levels.*

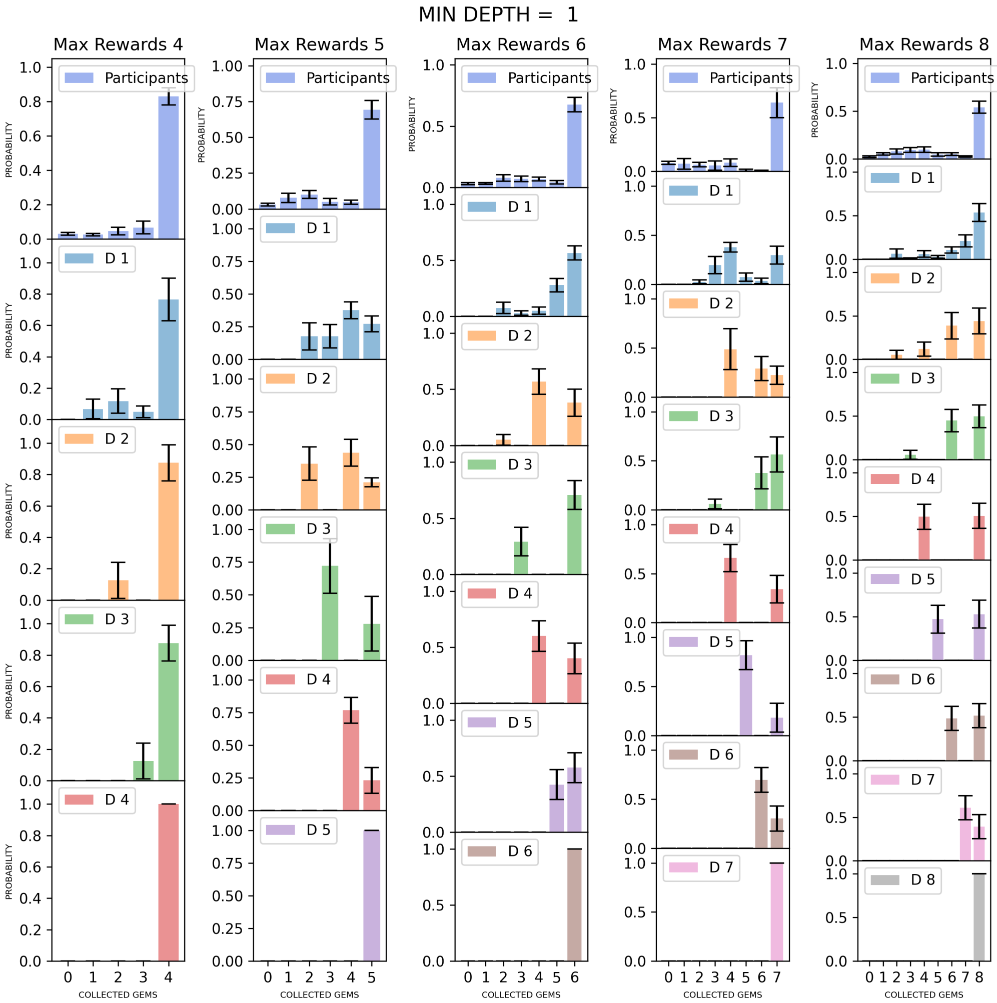


***Figure S5. Average distribution of gems collected by participants (first row) and by the planners D1-D8, which use planning depth 1 to 8, during the solution of problems requiring minimum depth of 1.*** *The panels show the results by splitting the problems by the maximum number of gems (Max Rewards). The plots show that for all problem classes, the 8 planning models collect different average gem distributions and are therefore distinguishable. Note that in this and the subsequent figures S6-S12, most of the planners fit well the peaks of the participants' distributions. Participants' data also show a variance for lower values of collected gems, which could be due to various sources of noise, not considered in our planners.*

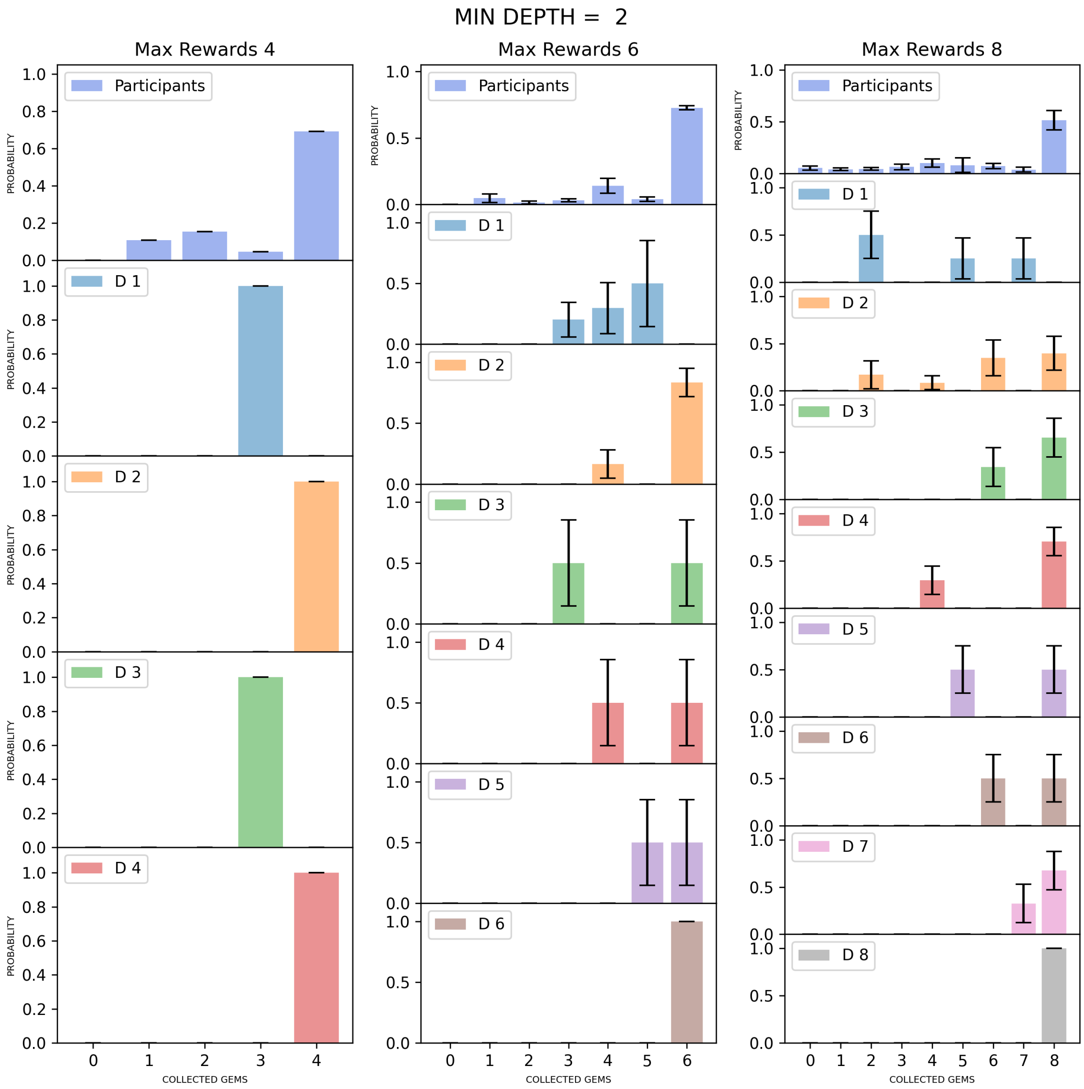


***Figure S6. Average distribution of gems collected by participants (first row) and by the planners D1-D8, which use planning depth 1 to 8, during the solution of problems requiring minimum depth of 2.*** *The panels show the results by splitting the problems by the maximum number of gems (Max Rewards). The plots show that for all problem classes, the 8 planning models collect different average gem distributions and are therefore distinguishable.*

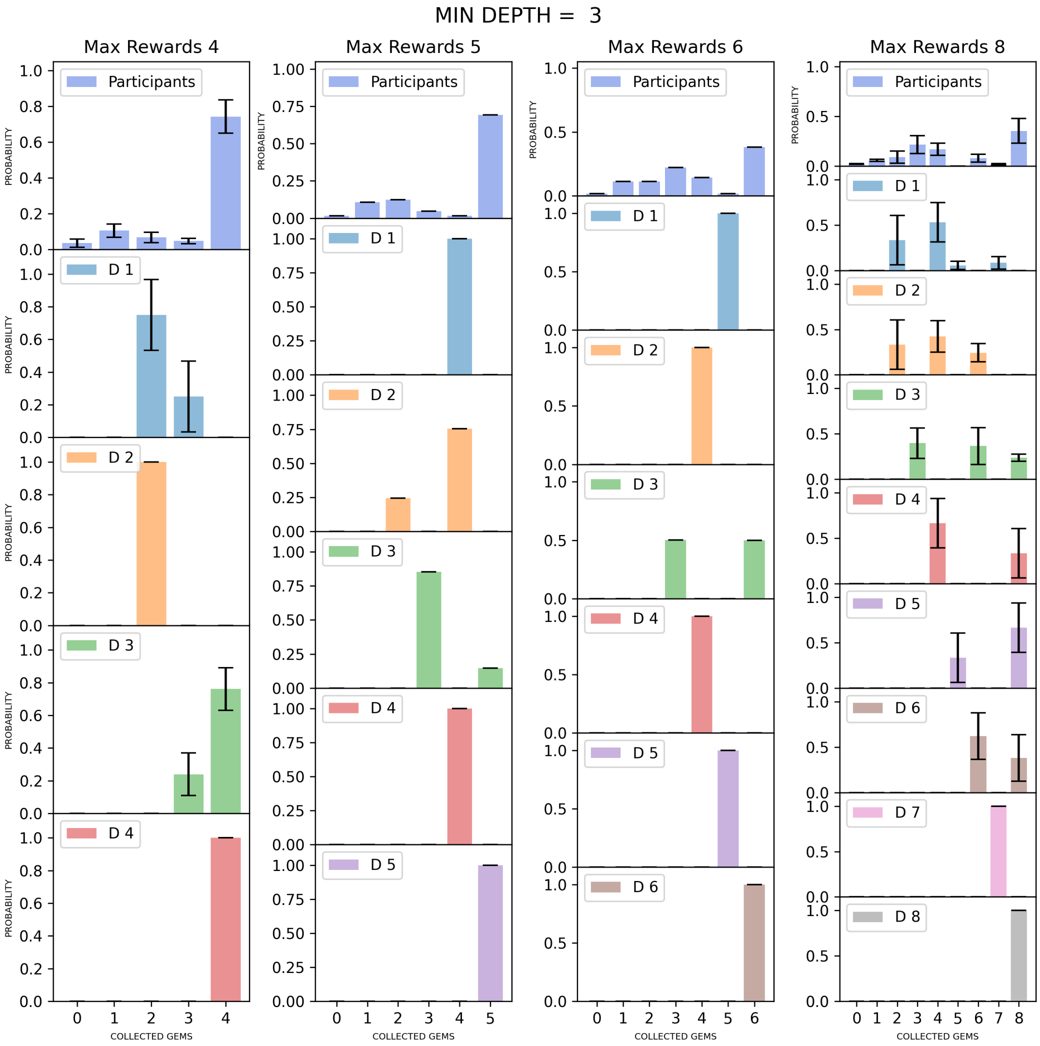


***Figure S7. Average distribution of gems collected by participants (first row) and by the planners D1-D8, which use planning depth 1 to 8, during the solution of problems requiring minimum depth of 3.*** *The panels show the results by splitting the problems by the maximum number of gems (Max Rewards). The plots show that for all problem classes, the 8 planning models collect different average gem distributions and are therefore distinguishable.*

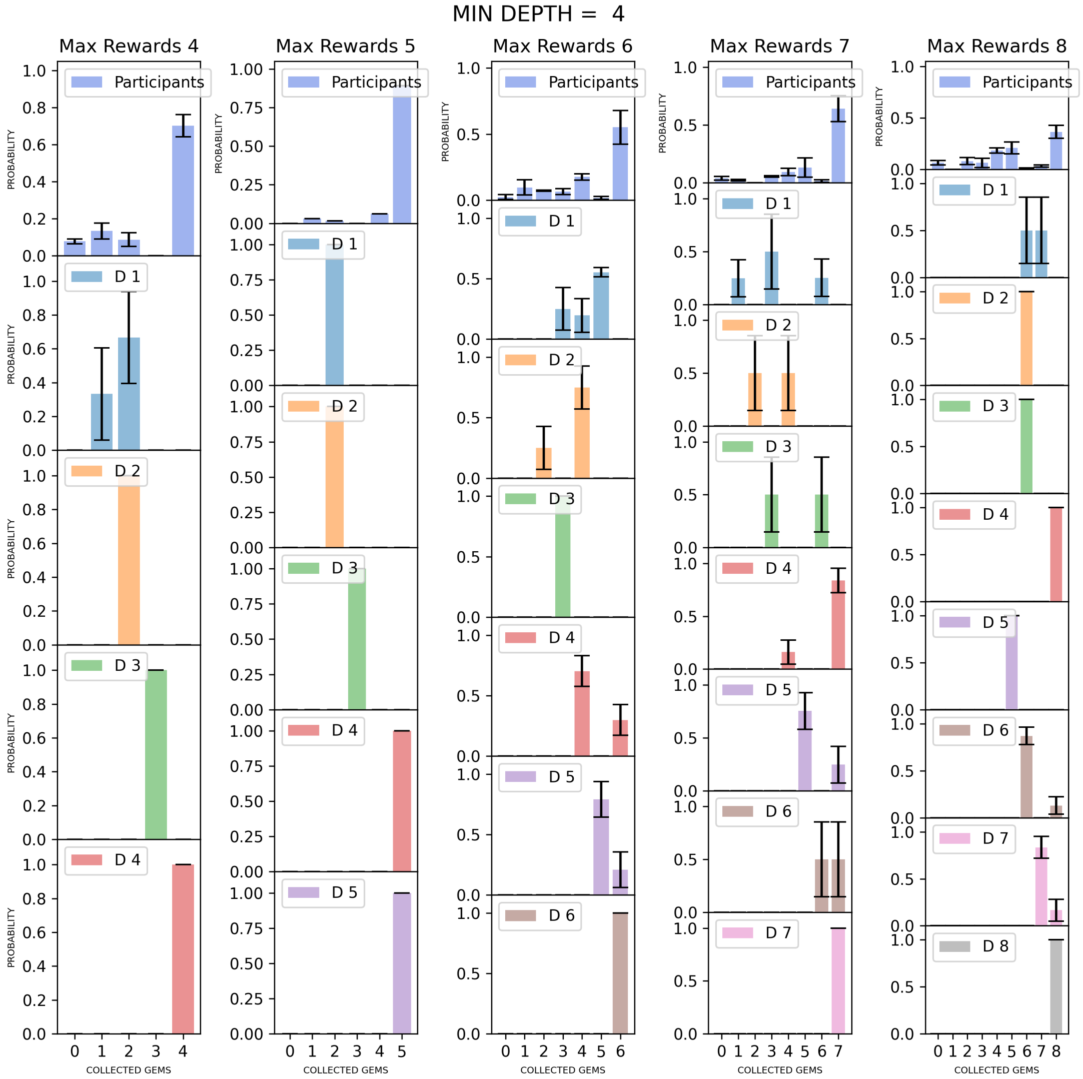


***Figure S8. Average distribution of gems collected by participants (first row) and by the planners D1-D8, which use planning depth 1 to 8, during the solution of problems requiring minimum depth of 4.*** *The panels show the results by splitting the problems by the maximum number of gems (Max Rewards). The plots show that for all problem classes, the 8 planning models collect different average gem distributions and are therefore distinguishable.*

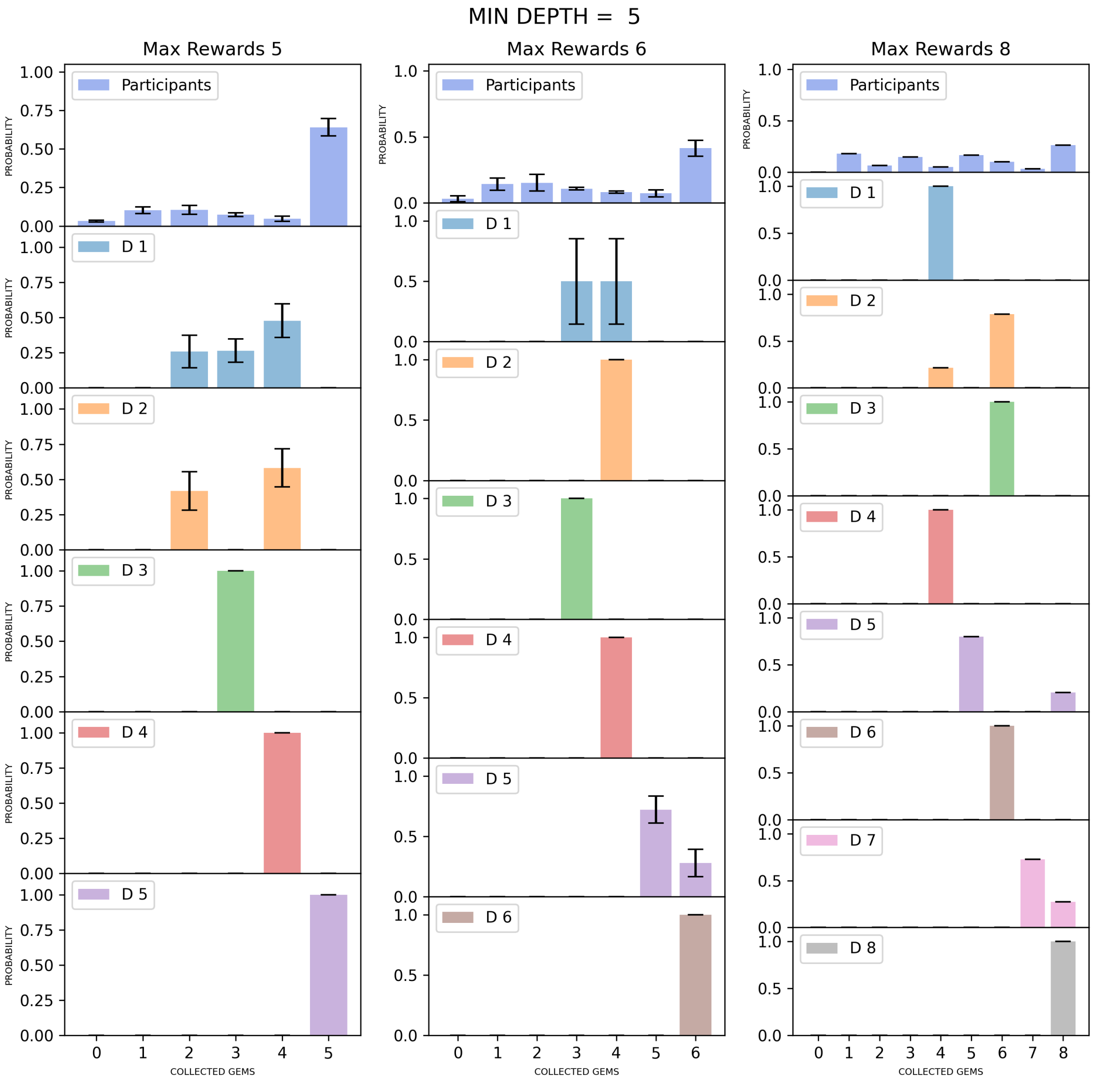


***Figure S9. Average distribution of gems collected by participants (first row) and by the planners D1-D8, which use planning depth 1 to 8, during the solution of problems requiring minimum depth of 5.*** *The panels show the results by splitting the problems by the maximum number of gems (Max Rewards). The plots show that for all problem classes, the 8 planning models collect different average gem distributions and are therefore distinguishable.*

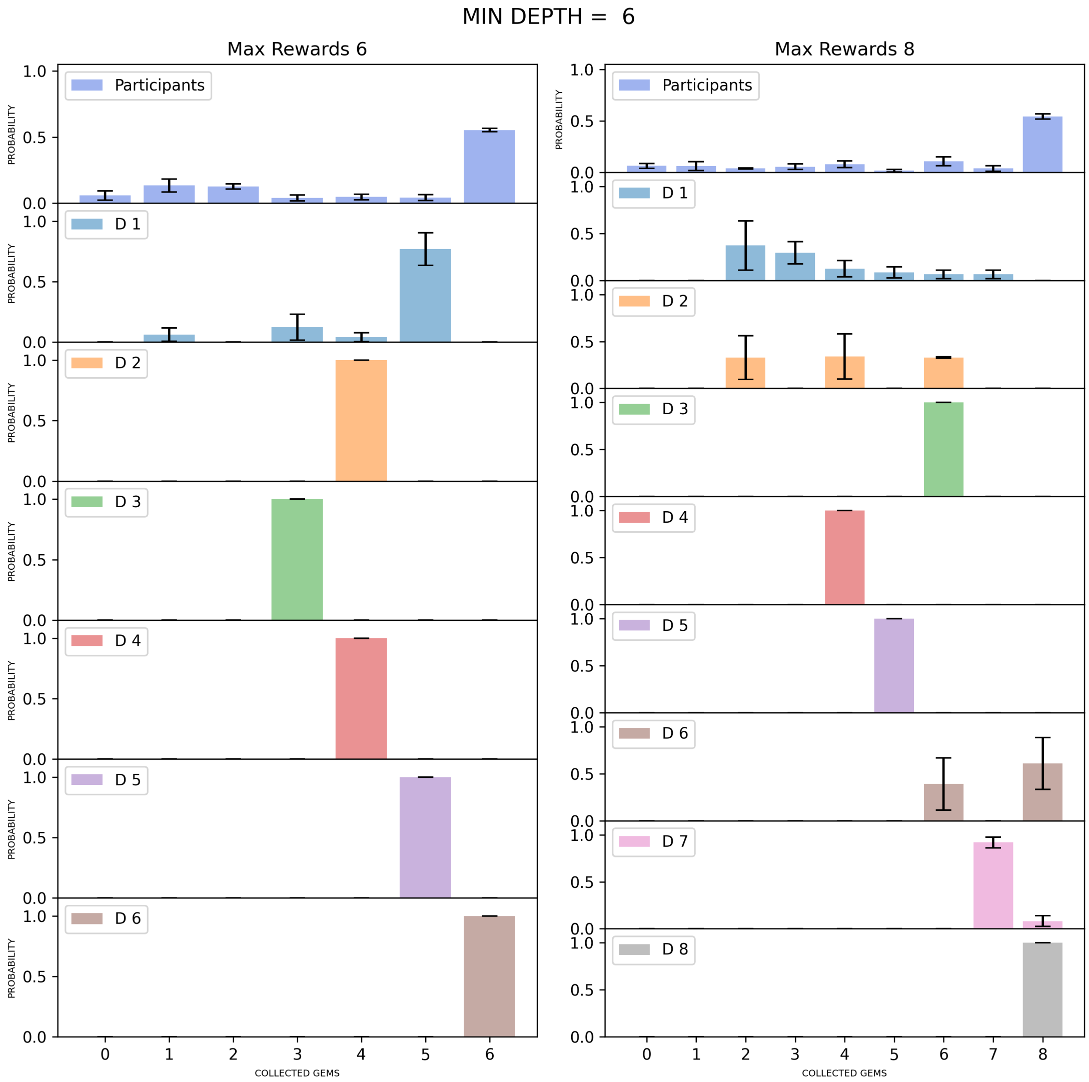


***Figure S10. Average distribution of gems collected by participants (first row) and by the planners D1-D8, which use planning depth 1 to 8, during the solution of problems requiring minimum depth of 6.*** *The panels show the results by splitting the problems by the maximum number of gems (Max Rewards). The plots show that for all problem classes, the 8 planning models collect different average gem distributions and are therefore distinguishable.*

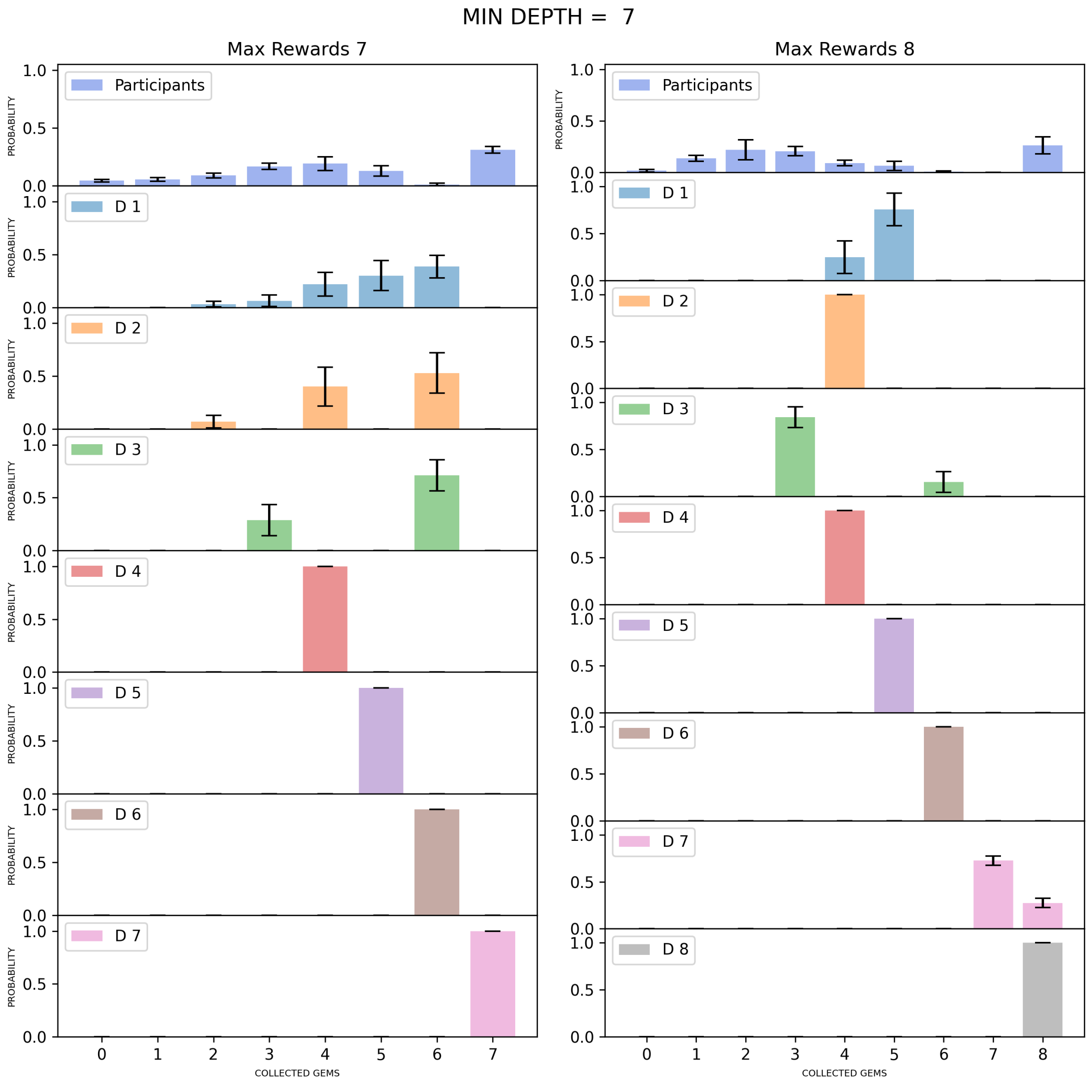


***Figure S11. Average distribution of gems collected by participants (first row) and by the planners D1-D8, which use planning depth 1 to 8, during the solution of problems requiring minimum depth of 7.*** *The panels show the results by splitting the problems by the maximum number of gems (Max Rewards). The plots show that for all problem classes, the 8 planning models collect different average gem distributions and are therefore distinguishable.*

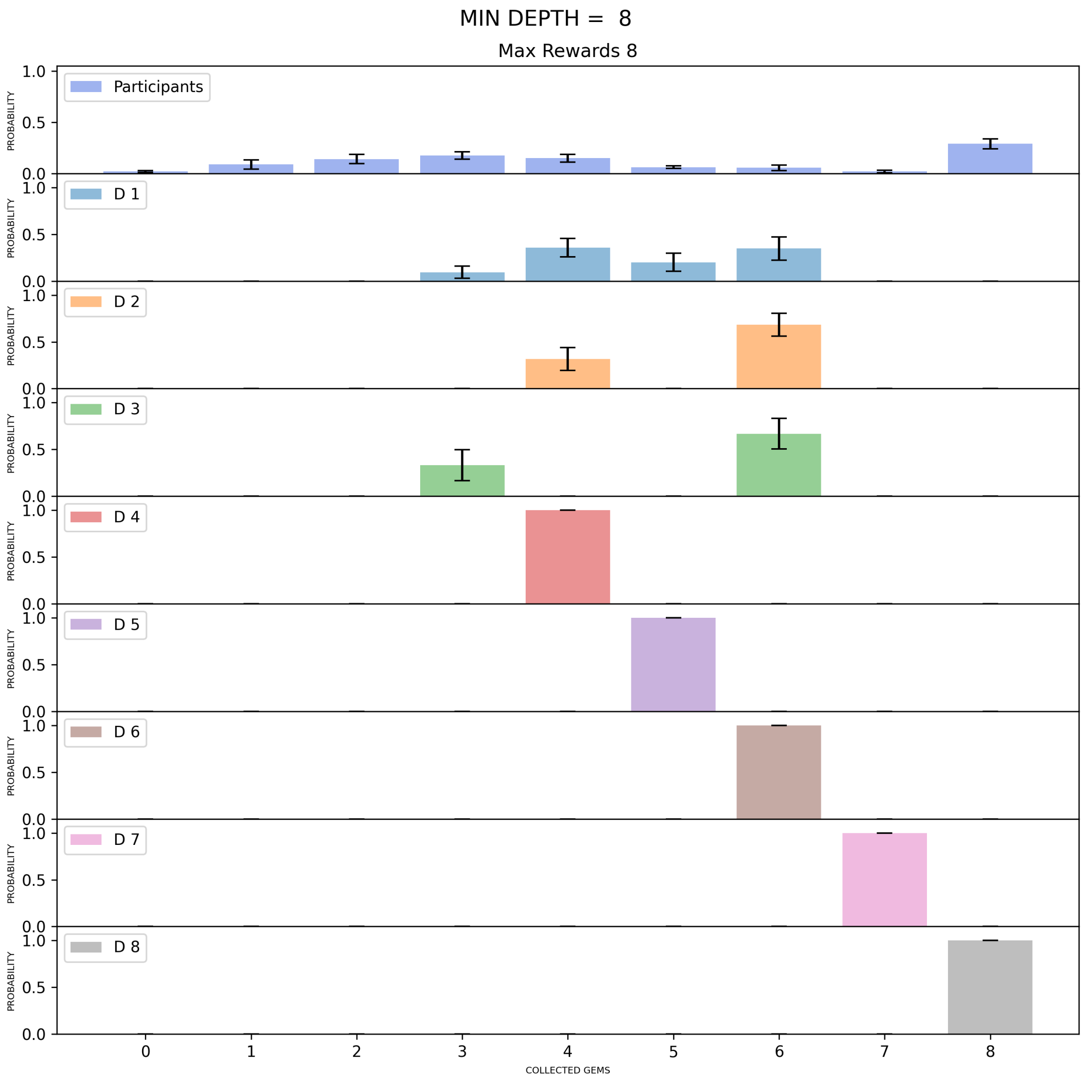


***Figure S12. Average distribution of gems collected by participants (first row) and by the planners D1-D8, which use planning depth 1 to 8, during the solution of problems requiring minimum depth of 8.*** *The panels show the results by splitting the problems by the maximum number of gems (Max Rewards). The plots show that for all problem classes, the 8 planning models collect different average gem distributions and are therefore distinguishable.*

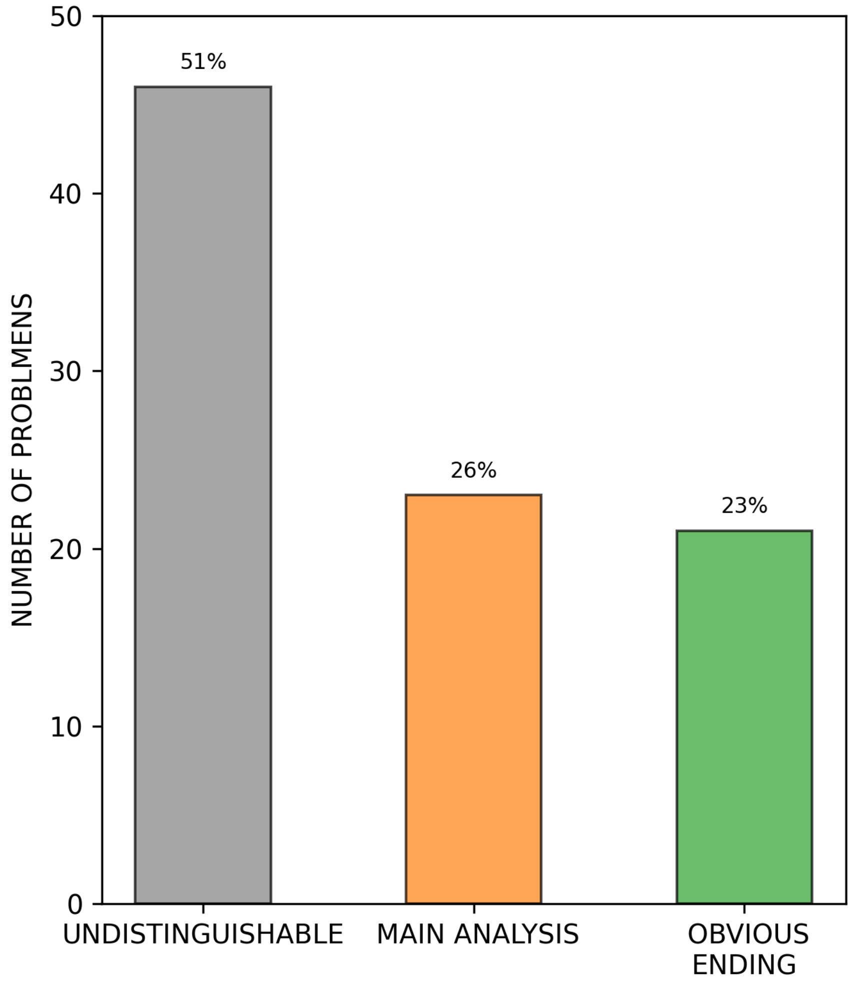


***Figure S13. Comparison of the performance of the planner used in the main analysis and of the "obvious ending" planner.*** *About half of the problems (51%, n = 46) do not have any obvious ending and hence cannot distinguish between the two models. The planner used in the main analysis and the "obvious ending" planner better explain participants' behaviour in 26% (n = 23) and 23% (n = 21) of the remaining problems, respectively.*

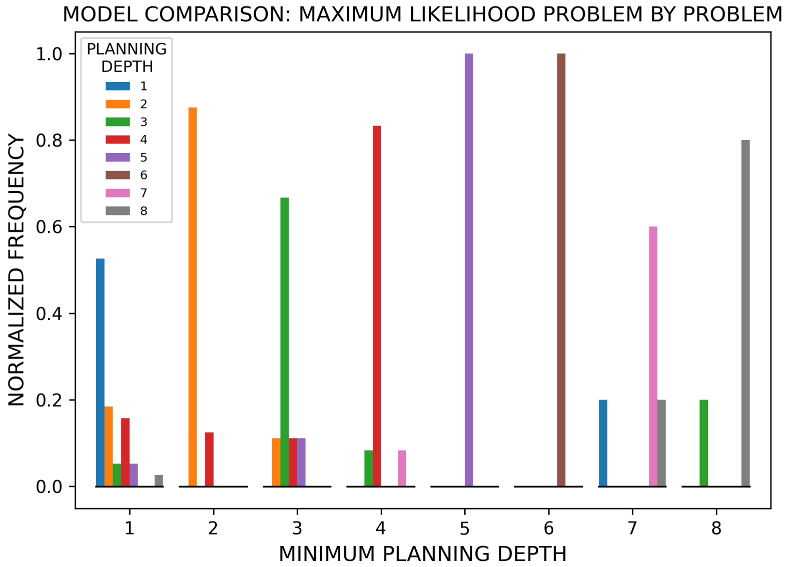


**Figure S14. *Comparison between participants and planning models, for the "obvious ending" planner.*** *The figure shows the number of times that each "obvious ending" planning model from depth 1 to 8 had the maximum likelihood of the gems collected before the first backtrack by the participants, in the problems of each of the 8 problem groups. Problems are grouped according to the minimum planning depth required to solve them, from 1 to 8, and color coded (see the legend).*

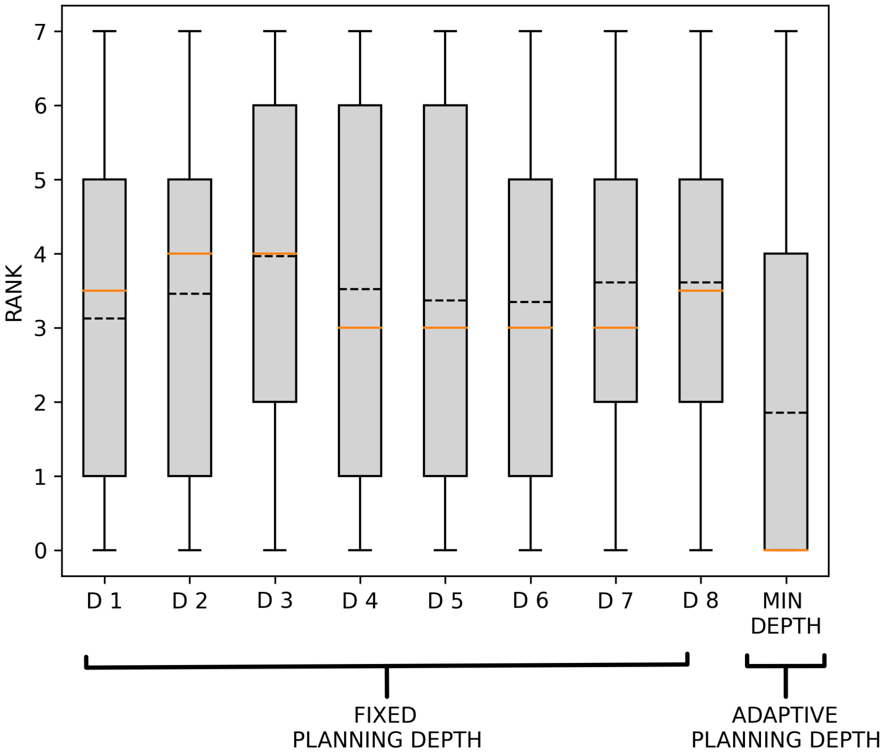


***Figure S15. Rank values across the 90 problems, for the "obvious ending" planner with fixed or adaptive planning depth****. The figure shows the distributions of ranks of planners using a fixed planning depth (D1-D8) across all the problems and of the planner using a planning depth adapted to the minimum required depth to solve a problem (Min depth). The black dotted lines show mean values, whereas the orange lines show median values. See the main text for explanation. The adaptive planning depth distribution is significantly smaller than any of the other distributions ($\chi^2(1, 90) = 31.51$, p-value $<5.0*10^{-5}$).*

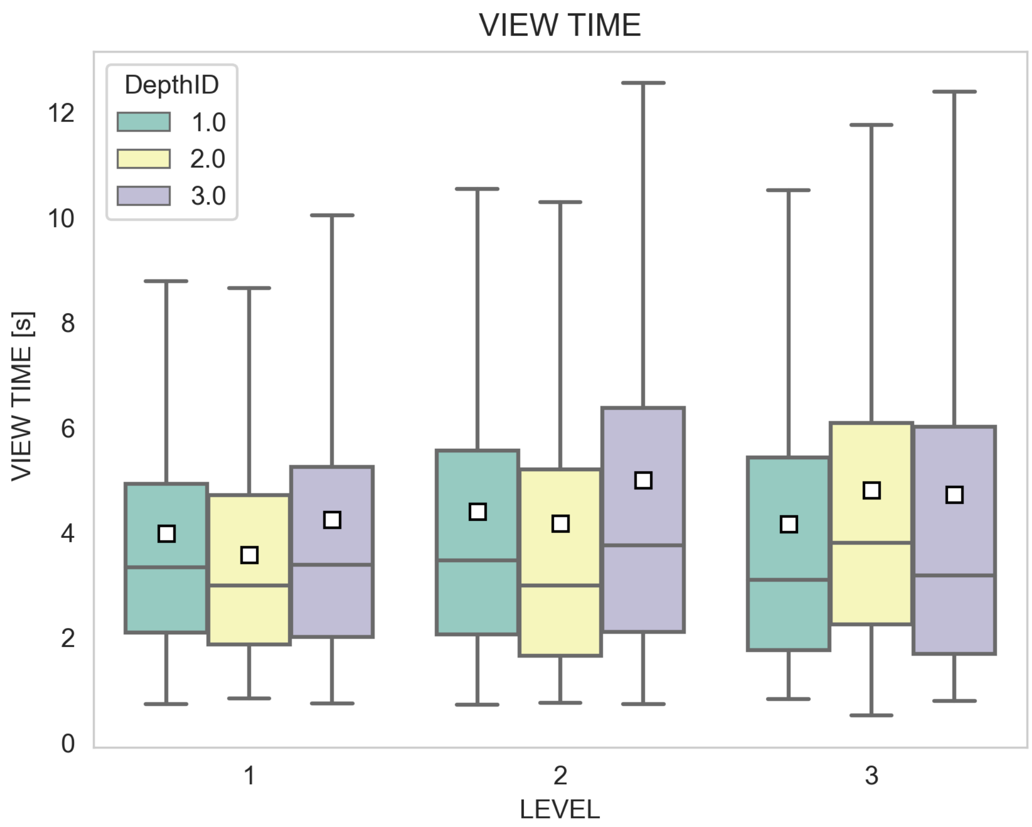


***Figure S16. View time (in seconds) as a function of the 3 levels of the experiment and of the 3 planning depths.*** *Results are organized by level and depth.*

***Table S1. View time regression.*** *A linear mixed-effect model is used to fit view time. Significance code (0< *** <0.001; 0.001** <0.01; 0.01 < * <0.05).*

| | **Regression coefficients** | **Std. error** | **DoF** | $\chi^2$ | **Pr(>$\chi^2$)** |
|---|---|---|---|---|---|
| **Intercept** | 3.59 | 0.39 | 1 | 68.24 | <$10^{-16}$ *** |
| **DepthID** | 0.076 | 0.12 | 1 | 0.37 | 0.54 |
| **Level** | 0.17 | 0.12 | 1 | 1.32 | 0.077 |
| **DepthID:Level** | 0.076 | 0.058 | 1 | 1.71 | 0.19 |

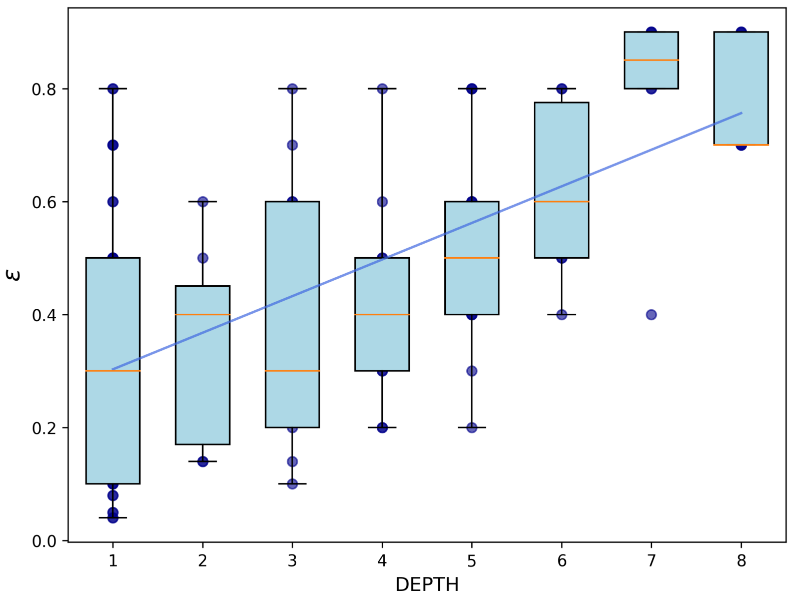


***Figure S17. Boxplot of the optimal ϵ and planning depth values as estimated via maximum likelihood.*** *The estimated parameters show a significant correlation ($p_{pearson}$ = 0.6, p-value < 10^-9). This means that in order to better describe participants performance, the model needs to use considerably higher noise values as the planning depth of the problem increases.*

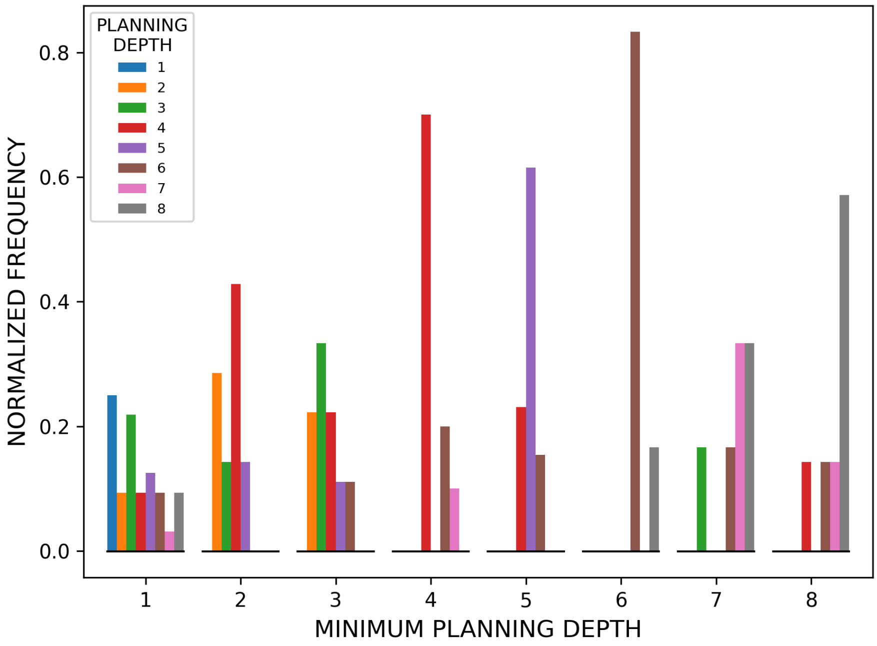


***Figure S18. Comparison between participants and planning models, for the for ϵ -greedy planner.*** *The figure shows the number of times that each ϵ -greedy planning model from depth 1 to 8 had the maximum likelihood of the gems collected before the first backtrack by the participants, in the problems of each of the 8 problem groups. Problems are grouped according to the minimum planning depth required to solve them, from 1 to 8, and color coded (see the legend).*

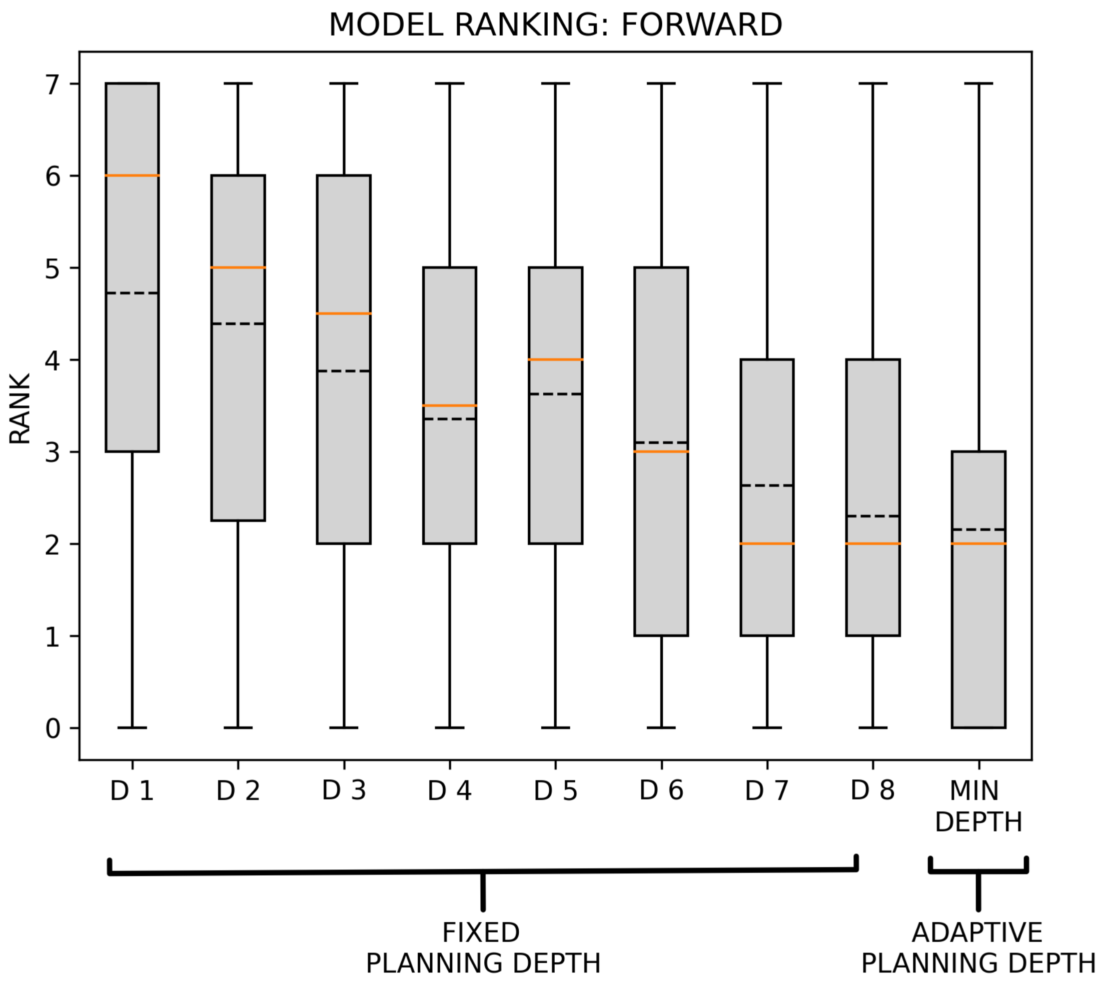


***Figure S19. Rank values across the 90 problems, for ϵ -greedy planner with fixed or adaptive planning depth**. The figure shows the distributions of ranks of planners using a fixed planning depth (D1-D8) across all the problems and of the planner using a planning depth adapted to the minimum required depth to solve a problem (Min depth). The black dotted lines show mean values, whereas the orange lines show median values. See the main text for explanation. The adaptive planning depth distribution is significantly smaller than the distributions of planning depth smaller than 7.*

**Pseudocode of the planning algorithms.**

The pseudocode of Algorithm 1 illustrates the procedure used by the planning model used in the main analysis to search a solution to the problems, at a fixed length. As explained in the main text, we used 8 variants of the same planning algorithm, which only differ for their planning depth, which ranges from 1 to 8. The algorithm selects the shortest path with a probability very close to 1; specifically, the probability of selecting the second-shortest path is of the order of $10^{-44}$. This is achieved by using a Softmax operator and by setting the precision (Beta) parameter to 100, hence closely approximating an Argmax operator.

Note that the paths from the current position to the next possible gems could not contain more gems than the depth considered at that step. For example, the paths obtained with depth = 1 could contain only one gem.

| **Algorithm1: Forward planning** | |
|---|---|
| **READ** *gems, max_ depth, start, graph* | |
| **SET** *full_path* = [], *collected_gems = []* | |
| **SET** *remaining_gems <- gems, current_node* <- *start* | |
| **WHILE** *remaining_gems:* | |
| | **SET** *depth* <- min(*remaining_gems, max_depth*) |
| | **CALCULATE** *all_simple_paths(current_position, remaining_gems, depth)* |
| | **IF** *all_simple_paths* ***is empty*** |
|  | **END** *"Reached a dead end!"* |
| | **ENDIF** |
| | **SAMPLE** *chosen_path <- softmax(all_simple_paths_lenghts, -BETA)* |
| | **UPDATE** *current_node, graph, remaining_gems, collected_gems* |
| | **CONCATENATE** *full_path <- chosen_path* |
| **ENDWHILE** | |
| **RETURN** *full_path* | |

The pseudocode of Algorithm 2 illustrates the procedure used by the **ϵ**-greedy planner.

```
Algorithm2: ϵ-greedy planner
READ  gems, max_ depth, start, graph, ϵ
SET full_path = [], collected_gems = []
SET  remaining_gems <- gems, current_node <- start
WHILE remaining_gems:
    SET depth <- min(remaining_gems, max_depth)
    IF rand < ϵ
        SAMPLE chosen_path<- current_node + random_neighboor
    ELSE
        CALCULATE all_simple_paths(current_position, remaining_gems, depth)
    IF all_simple_paths  is empty
        END "Reached a dead end!"
    ENDIF
    SAMPLE chosen_path <- softmax(all_simple_paths_lenghts, -BETA)
    UPDATE  current_node,  graph, remaining_gems, collected_gems
    CONCATENATE  full_path <- chosen_path
ENDWHILE
RETURN full_path
```
